# Trans-regenerational RNAi Memory in Planarians

**DOI:** 10.64898/2026.03.11.711021

**Authors:** Prakash V. Cherian, Idit Aviram, Uri Weill, Tamar Shapira, Sarit Anava, Hila Gingold, Jochen C. Rink, Oded Rechavi, Omri Wurtzel

**Affiliations:** School of Neurobiology, Biochemistry and Biophysics, George S. Wise Faculty of Life Sciences & Sagol School of Neuroscience, Tel Aviv University, Tel Aviv, Israel; Department of Tissue Dynamics and Regeneration, Max Planck Institute for Multidisciplinary Sciences, Göttingen, Germany; Faculty of Biology und Psychology, Georg-August-University Göttingen, Göttingen, Germany

**Author notes:** These authors contributed equally (Prakash V. Cherian, Idit Aviram). These authors contributed equally (Oded Rechavi, Omri Wurtzel).

## Abstract

How cells and organisms preserve molecular memory is a central question in biology. Chromatin states can maintain cellular memory in animals, yet DNA and histone modifications are typically reset in the germ line and early embryo. In *Caenorhabditis elegans*, RNA-dependent RNA polymerases (RdRPs) amplify small RNAs and enable transgenerational gene regulation. However, many organisms, including mammals, lack canonical RdRPs. Here we find that planarians, highly regenerative flatworms that lack canonical RdRPs and critical nuclear RNAi machinery, mount long-term heritable gene regulatory responses. We show that ingestion of double-stranded RNA induces potent, sequence-specific silencing that persists for months, withstands repeated cycles of whole-body regeneration, and can be transferred between animals. This long-term silencing, heritable during asexual reproduction, follows two phases: an early, transient response to exogenous RNA, and a stable, cell-autonomous memory phase that maintains repression long after the RNA trigger is gone. The memory phase is maintained post-transcriptionally, as transcription and chromatin accessibility remain unchanged while mature mRNA is depleted. This phase coincides with the emergence of A-tailed 21-23-nt antisense small RNAs and with silencing responses that act in *trans* and spread across the length of the target transcript. These results reveal a two-step, canonical RdRP-independent mechanism for persistent RNAi memory in a regenerating animal and establish planarians as a model for RNA-based inheritance, illuminating a novel mode by which RNA can encode durable gene-silencing states in metazoans.

## Introduction

For more than a century, the idea that acquired traits could be transmitted across generations has been met with skepticism, largely because it challenges the prevailing view that inheritance in animals is fully explained by DNA. This skepticism has been reinforced by the observation that in mammals chromatin modifications are extensively reprogrammed in the germ line and early embryo, effectively resetting the epigenetic landscape with each generation. Yet, not all animals necessarily conform to this paradigm.

Planarian flatworms can reproduce asexually by fission followed by regeneration, which bypasses the zygote and allows transgenerational continuity of adult tissues. This asexual reproduction is achieved using pluripotent stem cells called neoblasts, which rebuild all tissues, effectively bypassing the germline ^1,2^. Such unusual biology raises a compelling question: can molecular states be stably maintained between fragments of the same body? The possibility that planarians might transmit information independently of DNA once fueled some of the most controversial experiments in memory research. In the 1960s, James V. McConnell reported that learned behaviors could survive decapitation and even be transferred to naïve worms through feeding ^3^, an idea later revisited with modern behavioral assays but still unexplained at the molecular level ^4^.

RNA interference (RNAi) constitutes a molecular mechanism that potentially enables this phenotypic persistence. This pathway is highly potent in diverse planarian model species (e.g., *S. mediterranea* and *D. japonica*). Planarian RNAi can be induced by feeding or injecting double-stranded RNA ^5,6^, and is the standard approach for gene silencing, as the ability to delete or edit genes has not been achieved yet in planarians. In many systems, RNA-guided silencing is transient and decays once the initial trigger is removed ^7,8^. Long-lasting or heritable silencing typically requires RNA-dependent RNA polymerases (RdRPs), which amplify small interfering RNAs from target transcripts, as seen in nematodes ^9–11^ and fungi ^12^. However, planarians lack recognizable eukaryotic RdRPs and key effectors required for systemic or heritable RNAi, such as RSD-2, NRDE-3, and HRDE-1 ^13^. They also likely lack DNA methylation ^14^, and therefore this mechanism cannot reinforce silencing RNAs as it does in plants (RNA-directed DNA methylation) ^15^. This raises a profound question: How can planarians mount potent and long-lasting RNAi-responses in the absence of known memory mechanisms?

Here, we systematically characterized the RNAi response in planarians and reveal that exposure to double-stranded RNA triggers silencing that endures for months and persists through multiple rounds of complete regeneration. Using transplantation and clonal analyses, as well as RNA-reporter assays, we show that the response unfolds in two distinct phases: an initial, transient gene silencing driven by the short-lived exogenous dsRNA, and a prolonged, cell-autonomous memory phase that maintains post-transcriptional gene suppression independently of exogenous dsRNA input. These findings uncover an unexpected, RdRP-independent mechanism for stable RNAi memory in a regenerating animal, expanding current models of RNA-based inheritance beyond polymerase and chromatin-centered paradigms. This discovery establishes a foundation for investigating alternative mechanisms of RNA-based inheritance in animals, and identifying their contributions to physiology.

## Results

### RNAi-induced phenotypes persist across multiple regeneration cycles in planarians

Planarians are known to mount a robust and organism-wide RNAi response and anecdotal evidence indicates that the gene silencing effects are particularly long-lasting ^16^. However, the duration of the RNAi response has not been systematically quantified so far. Towards this goal, we selected two genes with strong and easily tractable RNAi phenotypes: (i) *ovo-1*, a transcription factor (TF) that is specifically expressed in a subset of neoblasts and eye progenitors and leads to the striking loss of the eyes when suppressed ^17^; and (ii) *ꞵ-catenin-1*, which is the mediator of canonical Wnt signaling in planarians and enforces the regeneration of heads in suppressed injured animals ^18–20^. Feeding *in vitro* synthesized dsRNA reproduced the expected phenotypes: head regeneration without eyes in *ovo-1* (RNAi) animals (Fig. 1A) and ectopic head formation following *ꞵ-catenin-1* inhibition (Fig. 1B).

**Figure 1.**
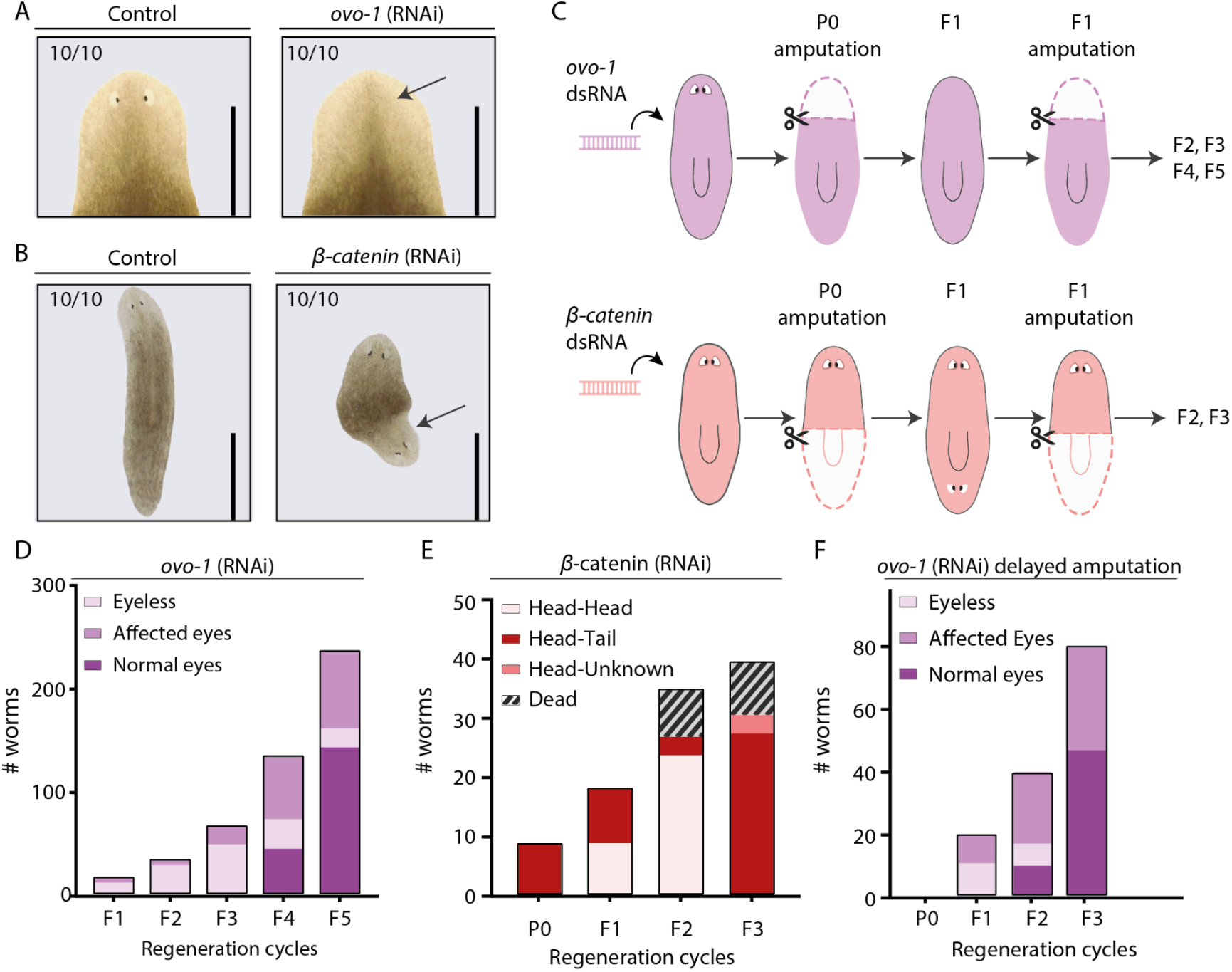
RNAi-induced phenotypes persist across successive regenerative cycles in planarians. (A) Representative images of *S. polychroa* 14 days post-amputation (dpa) following 12 feedings of *ovo-1* or control dsRNA. Arrow indicates the loss of eyes in the *ovo-1* (RNAi) animal. (B) Representative images of *S. mediterranea* 14 dpa following two feedings of *ꞵ-catenin* or control dsRNA. The arrow indicates ectopic head formation. Matching controls for both assays exhibited normal phenotypes (n = 10 per group). Experiments were repeated at least three times with >10 animals per experimental group. Scale bars = 1 mm. (C) Schematic detailing the experimental timeline for successive cycles of amputation and regeneration. (D) Quantification of eye defects in *ovo-1* (RNAi) animals at each regeneration cycle. Eye defects persisted through the fifth regeneration cycle (F5) (n = 295 total; 108/295 exhibiting defects). (E) Quantification of abnormal anterior-posterior axis formation in *ꞵ-catenin* (RNAi) animals. Ectopic head regeneration persisted across three regeneration cycles (F1-F3). (F) Quantification of eye defects in *ovo-1* (RNAi) animals subjected to delayed amputation. Animals were amputated at 5 weeks post-feeding instead of 1 week (Fig. S1A) to isolate the effect of absolute time elapsed from the extent of cell turnover. Approximately 45% of animals exhibited eye development defects at the final measured cycle (34/80, n = 80), indicating that absolute time elapsed strongly dictates the persistence of the RNAi effect.

Next, we assessed the duration of the RNAi response by measuring phenotype penetrance across successive rounds of amputation and regeneration in two planarian species, the asexual strain of *Schmidtea mediterranea* and a sexually reproducing strain of its sister species *Schmidtea polychroa* (Fig. 1C). Animals were fed dsRNA targeting *ovo-1* or *ꞵ-catenin-1* and then subjected to repeated cycles of amputation and regeneration without further dsRNA exposure to monitor the persistence of the induced phenotypes. Remarkably, *ovo-1* silencing, evident as eye development defects, was maintained throughout five consecutive regeneration cycles in 36% of the animals, despite the extensive cell turnover accompanying each cycle ^21,22^. Thus, more than one-third of the treated animals retained the *ovo-1* (RNAi) phenotype for the full duration of the three-month experiment (Fig. 1D). Similarly, *ꞵ-catenin-1* silencing led to the regeneration of ectopic heads, a phenotype that was maintained across several regeneration cycles. However, animals showing these severe patterning defects rarely survived beyond the third cycle (Fig. 1E).

We next assessed whether the persistence of the RNAi phenotype was influenced by the extent of cell turnover. To test this, we introduced a delay after the final dsRNA feeding before amputation, resulting in fewer amputations within the same overall time window. After completing dsRNA feeding, one group of animals was amputated one week later, whereas another group remained intact for five weeks before amputation (Fig. S1A). The strength of the RNAi response, assessed by phenotype persistence (Fig. S1B), was unaffected by the initial delay before the first amputation, with both groups showing similar outcomes once they were amputated and regenerated in parallel. Instead, phenotype persistence correlated primarily with the time elapsed since the final dsRNA feeding (Fig. 1F). These results indicated that dsRNA exposure induced a long-lasting RNAi state that remained stable over time, independent of injury or repeated tissue regeneration.

### Exogenous dsRNA is short-lived

The RNAi-induced phenotypes lasted multiple regeneration cycles. One possible explanation is that “leftover” dsRNA from the parent persists in the offspring, and despite its expected dilution, it remains in quantities sufficient for inducing silencing. We therefore developed a qPCR-based assay to measure the concentration of exogenous dsRNA over time via the amplification of a synthetic adapter sequence present only in the exogenous dsRNA (Fig. 2A; Methods). We found that dsRNA levels declined by approximately 1000-fold within one week following dsRNA ingestion and were essentially undetectable (10^4^-fold reduction) four weeks after uptake (Fig. 2B). Furthermore, analysis of dsRNA integrity revealed that while the silencing effect lasts multiple months, only ∼1% of the detectable dsRNA remained full-length at 7 and 14 days post feeding (dpf), indicating that the dsRNA was rapidly degraded or processed (Fig. 2C; Methods).

**Figure 2.**
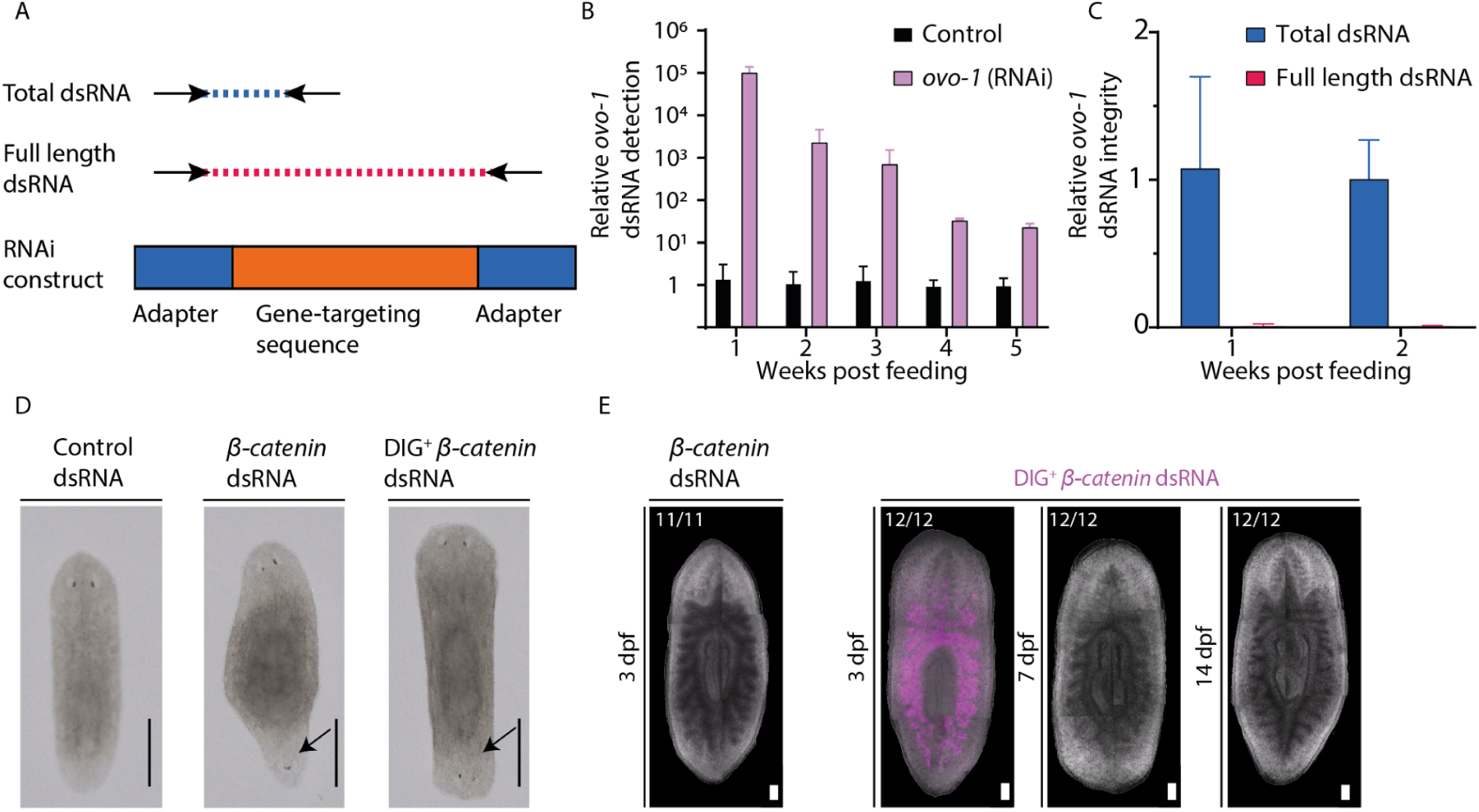
Exogenous dsRNA depletes rapidly following ingestion. (A) Schematic of the qPCR strategy designed to selectively amplify total versus full-length exogenous dsRNA. (B) qPCR quantification of relative *ovo-1* dsRNA levels in homeostatic *ovo-1* (RNAi) and control animals. Exogenous *ovo-1* dsRNA abundance decreases by orders of magnitude across the 5 weeks following the final feeding. Data represent mean ± standard deviation. (C) Quantification of total versus full-length *ovo-1* dsRNA at 1 and 2 weeks post-feeding. The near-complete absence of full-length dsRNA indicates the initial trigger undergoes rapid processing or degradation. Data represent mean ± standard deviation. (D) Representative images of regenerated animals following two feedings of control, *β-catenin*, or DIG-labeled *β-catenin* dsRNA. Animals fed DIG-labeled dsRNA successfully regenerate ectopic heads (arrows), confirming that DIG labeling does not impair RNAi efficacy. The fraction of animals displaying the phenotype is indicated in each panel. Scale = 0.5 mm. (E) *In situ* detection of DIG-labeled *β-catenin* dsRNA at 3, 7, and 14 days post-feeding (dpf). Exogenous dsRNA strongly labeled the intestine at 3 dpf but was largely depleted by 7 dpf. Unlabeled *β-catenin* dsRNA (3 dpf) serves as a negative control for labeling specificity (Methods). Scale = 100 µm. The fraction of animals displaying the representative pattern is indicated in each panel.

To track the fate of the dsRNA in the animal, we fed worms with *ꞵ-catenin* dsRNA labeled with digoxigenin (DIG) detectable by anti-DIG antibodies (Methods). All amputated RNAi animals displayed the expected phenotypes (Fig. 2D), confirming that DIG-labeling did not impair the biological activity or systemic spreading of the RNAi response. *In situ* detection of the DIG-labeled dsRNA showed that it was initially concentrated in the planarian intestine but was largely depleted from this region by seven days following dsRNA ingestion (Fig. 2E). This rapid depletion of dsRNA, in line with previous *in situ* hybridization studies ^6^, indicated that the maintenance of the RNAi phenotype did not rely on the continued presence of dsRNA. Indeed, when we examined dsRNA concentration inside cells, by purifying planarian cells 6 and 14 days following dsRNA ingestion using fluorescence-activated cell sorting (FACS), we did not detect appreciable levels of exogenous dsRNA (Fig. S2A-B; ∼10^4^-fold less than in whole tissue RNA extractions). Therefore, planarian cells were unlikely to be a long-term reservoir of unprocessed dsRNA. In summary, the rapid decay of dsRNA contrasts sharply with the long-lived silencing response, implying that RNAi maintenance depends on memory-like mechanisms rather than on persistent dsRNA.

### Tissue transplantations reveal a critical window for RNAi memory activation prior to stable maintenance

We observed a rapid decline in dsRNA concentration following ingestion, prompting us to consider two mechanisms for continued systemic spreading: (i) spreading continues, even though the dsRNA is barely detectable, via some other secondary effector (not dsRNA), (ii) systemic RNAi spreading stops, because it depends on higher levels of dsRNA. To distinguish between the two possibilities, we used a tissue transplantation assay and exchanged tissues between RNAi-treated hosts and naive donors. In this experiment, we fed animals dsRNA targeting *ovo-1*, and then irradiated the animals in order to ablate their neoblast population (Methods). Next, we transplanted tissues isolated from naive unirradiated donor animals into the irradiated hosts (Fig. 3A). We assessed whether the RNAi effect was transferred to the grafted donor tissues. If the RNAi effect was transferable, the graft neoblasts would fail to produce eye cells, despite being able to regenerate other tissues; if not, the graft neoblasts would regenerate all missing structures, including eyes. Notably, all grafts (n = 18) transplanted one week after dsRNA ingestion regenerated head structures. However, 83% produced fully eyeless heads (n = 18, eyeless = 15/18), and the remaining grafts showed partial eye defects (Fig. 3B), indicating that the RNAi effect was transferred from the host to the graft neoblasts. A similar transfer of the RNAi effect from RNAi host to naive graft, one week following dsRNA ingestion, was observed when we suppressed a different gene, *fir-1* ^23^ (n = 7/7; Fig. S3A-B).

**Figure 3.**
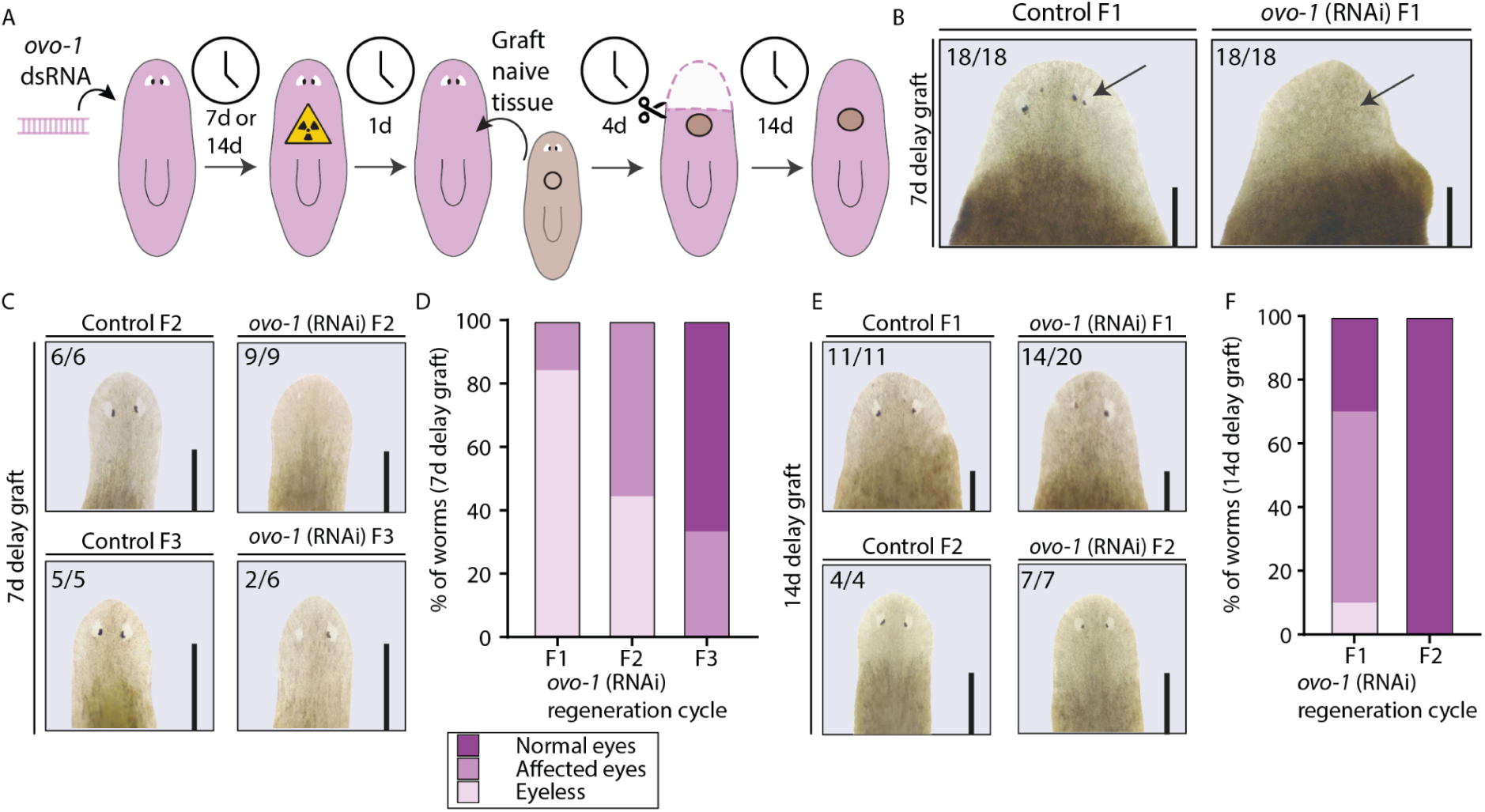
RNAi-mediated gene silencing can spread horizontally within a critical time window. (A) Schematic of the tissue grafting experiment. *ovo-1* (RNAi) and control animals were starved for 7 or 14 days following the final dsRNA feeding (Methods), then lethally irradiated to ablate neoblasts. Naive, unirradiated tissue was grafted onto the hosts 24 hours post-irradiation. After 4 days of graft integration, heads were amputated and the regenerated tissue was assessed 14 days later. (B, C) Representative images of regenerated heads from animals grafted 7 days post-RNAi. Panel B shows the first regeneration cycle (F1), where arrows indicate normal eyes in the control and absent eyes in the *ovo-1* (RNAi) host. Panel C shows subsequent cycles (F2, F3). (D) Quantification of eye phenotypes following the 7-day delay graft. The silencing effect successfully transferred to the naive tissue and persisted into F3. Data represent the percentage of *ovo-1* (RNAi) animals exhibiting each phenotype (n = 18, 9, and 6 for F1, F2, and F3, respectively). All matching control grafts regenerated normal eyes (n = 18, 6, 5). (E) Representative images of regenerated heads from animals grafted 14 days post-RNAi across the F1 and F2 cycles. Scale bars = 1 mm. (F) Quantification of eye phenotypes following the 14-day delay graft. Phenotypic transfer to the naive tissue was weaker than in the 7-day delay cohort and fully resolved by F2 (*ovo-1* n = 20 and 7 for F1 and F2, respectively). All matching control grafts regenerated normal eyes (n = 11, 4).

We next tested whether the RNAi effect remained transferable after dsRNA levels declined. When grafts were transplanted two weeks after *ovo-1* dsRNA feeding, only 10% of animals regenerated fully eyeless heads, indicating a much milder *ovo-1* suppression phenotype (Fig. 3C). This identifies a critical one-week window during which dsRNA, or its effectors, can be transferred to grafted neoblasts, consistent with dsRNA levels having dropped to roughly 1% of their one-week post-feeding concentration (Fig. 2B).

We further examined whether the *ovo-1* RNAi phenotype, established in grafts transplanted one week post-dsRNA ingestion, persisted through subsequent regeneration cycles. In all animals transplanted one week following the RNAi feedings (n = 18), the phenotype endured through three cycles of regeneration, suggesting that once established, RNAi memory is both robust and long-lasting (Fig. 3C-D). Conversely, far fewer grafts transplanted two weeks after *ovo-1* dsRNA feedings developed a phenotype (Fig. 3E-F). Moreover, the transient phenotype failed to persist in subsequent regeneration cycles (n = 7/7), which indicated that RNAi-mediating effectors are required to establish long-term RNAi memory.

Together, these experiments demonstrate that long-term RNAi persistence occurs in two distinct phases. Phase 1 is a systemic, non-autonomous period where exogenous dsRNA spreads to initiate silencing, a process limited by the short half-life of dsRNA or its processing products ^24^. Phase 2 represents a memory phase where cells maintain RNAi independently of dsRNA. This durable, cell-autonomous, and heritable silencing forms only in cells exposed to RNAi effectors during Phase 1. This transition shifts RNAi from a transient systemic input to a stable memory. We next investigated the molecular mechanisms underlying this process.

### Transcription is not suppressed during RNAi memory in planarians

In diverse organisms, RNAi memory is maintained via *cis*-acting mechanisms leading to transcriptional repression at the targeted locus ^15,25,26^. To determine whether RNAi memory in planarians operates through transcriptional repression as well, we measured both pre-mRNA and mature mRNA expression levels at multiple time points following dsRNA treatment using a qPCR-based detection strategy (Fig 4A; Methods). This dual approach allowed us to differentiate between transcriptional and post-transcriptional silencing. Specifically, if transcription were suppressed at the targeted locus, both pre-mRNA and mature mRNA levels would decline. However, if only the mature mRNA level were reduced, it would suggest that transcription was unaltered and that silencing occurred post-transcriptionally.

**Figure 4.**
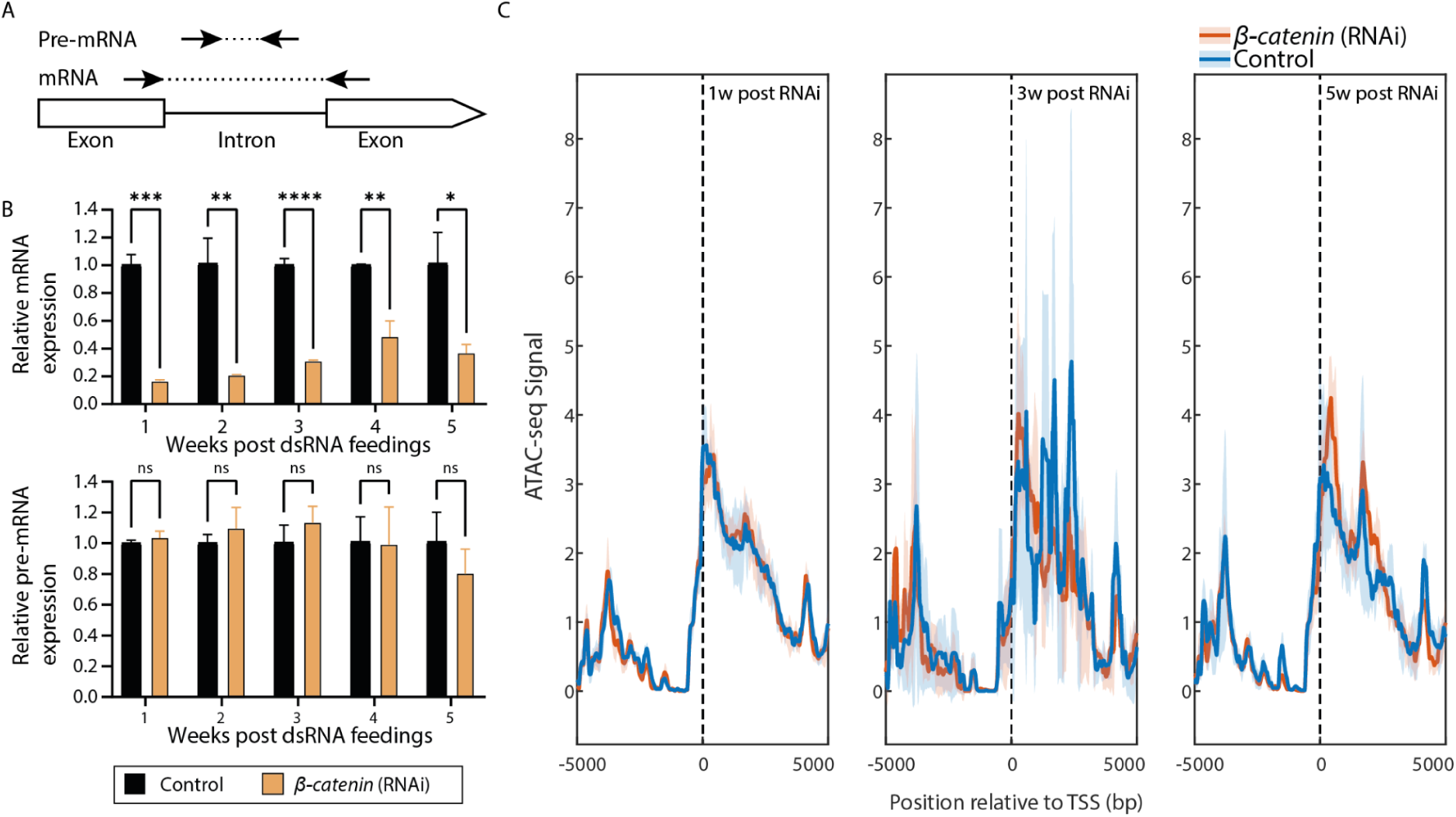
RNAi-mediated gene silencing does not involve transcription suppression in *S. mediterranea*. (A) Schematic of the qPCR primer design (Methods) used to distinguish pre-mRNA (intron-spanning) from mature mRNA (exon-spanning). (B) qPCR quantification of mature *ꞵ-catenin* mRNA (top) and pre-mRNA (bottom) relative to controls. Mature mRNA levels decrease significantly, whereas pre-mRNA levels remain stable over the 5-week time course. Data represent mean ± standard deviation. Statistical significance was determined using multiple t-tests with Holm–Šídák correction (*P < 0.05, **P < 0.01, ***P < 0.001, ****P < 0.0001; ns = not significant). (C) ATAC-seq profiles of chromatin accessibility relative to the transcription start site (TSS = 0) at 1, 3, and 5 weeks post-dsRNA ingestion. The accessibility signal shows no difference between *ꞵ-catenin* (RNAi) and control animals at any time point. Shown is the mean (solid color), and standard deviation (light color).

To distinguish between these alternatives, we administered 2 feedings of *ꞵ-catenin* dsRNA to intact (non-regenerating) planarians (Methods), and monitored gene expression over five weeks. We used two sets of primer pairs targeting a region that does not overlap the *ꞵ-catenin* dsRNA sequence (Fig. 4A; Methods). One primer pair, designed to amplify mature mRNA across an exon–exon junction, selectively quantified mature transcripts, while a second primer pair, designed to anneal to an intron sequence, reported on pre-mRNA levels (Fig 4A; Methods). In agreement with our previous findings, *ꞵ-catenin* dsRNA levels were already virtually undetectable as early as 1 week post-RNAi (Fig. 2E-F), confirming our previous conclusion that the RNAi response persists beyond the presence of exogenous dsRNA. As expected, the mature *ꞵ-catenin* mRNA levels were depleted throughout the experiment (Fig. 4B), further confirming that prolonged gene silencing does not necessitate the continued presence of dsRNA. Strikingly, pre-mRNA levels remained unchanged following RNAi treatment at all time points (Fig. 4B; Methods).

To further test whether the *ꞵ-catenin* locus remained transcriptionally active during the memory phase of the RNAi response, we assayed genome-wide chromatin accessibility using ATAC-seq ^27^ at 1, 3, and 5 weeks following dsRNA ingestion. Across these datasets, we identified 19,428 consensus peaks, enabling the detection of putative promoter regions (Table S1; Methods). Using this framework, we identified a putative *ꞵ-catenin* promoter region upstream of its transcription start site (Fig. S4). Strikingly, chromatin accessibility at both the *ꞵ-catenin* promoter and gene body remained unchanged across all time points and was indistinguishable between *ꞵ-catenin* (RNAi) and control animals (Fig. 4C). Together, the qPCR and ATAC-seq data show that RNAi memory does not involve transcriptional repression or chromatin remodeling at the targeted locus, strongly indicating that the memory phase of the planarian RNAi response likely operates entirely post-transcriptionally.

### No evidence for canonical RdRP-mediated amplification of sRNAs in planarians

A prominent mechanism known to sustain RNAi memory in other organisms relies on RNA-dependent RNA polymerase (RdRP), which generates secondary antisense small RNAs (sRNAs) to amplify and maintain the silencing response ^9,28,29^. Thus, we searched for putative RdRP-encoding sequences in five planarian genomes ^30^ using multiple approaches, including sequence-similarity analysis and protein domain predictions (Methods). We did not find any sequence resembling eukaryotic RdRPs in the planarian genomes. By contrast, when these analyses were performed with the *Macrostomum lignano* genome, a non-planarian flatworm that is also able to mount a robust RNAi response ^31^, we readily identified multiple candidates for eukaryotic-type RdRPs (Methods). However, the possibility remained that atypical or non-eukaryotic RdRPs might nevertheless generate secondary sRNAs in planarians.

To detect potential RdRP-derived secondary sRNAs, we sequenced small RNAs from naive planarians with and without tobacco acid pyrophosphatase (TAP) treatment. TAP converts 5’-triphosphates, a characteristic of RdRP products in C. elegans ^9,29^, into illumina sequencing-ready 5’-monophosphates. Analysis of the whole genome revealed two distinct sRNA populations in both libraries: 20–23 nt miRNAs^32^ and 32–34 nt piRNAs ^33^ (Fig. S5A-D). The 32–34 nt population exhibited canonical piRNA features, including a 5’ uridine bias and a +10 adenosine (Fig. S5E). Crucially, TAP treatment yielded no differences in sRNA population size or sequence characteristics compared to untreated samples (Fig. S5A-D). This absence of TAP-dependent enrichment indicates that naive planarians lack the canonical 5’-triphosphate signature associated with RdRP amplification.

We next tested whether ingestion of dsRNA triggered sRNA amplification at the targeted gene locus. Planarians were fed dsRNA targeting *ꞵ-catenin*, and seven days post-feeding we isolated sRNAs and used TAP treatment to distinguish between primary and putative secondary sRNAs (Methods). Following sequencing of the TAP-treated and untreated samples, we examined mapping of sRNAs at the *ꞵ-catenin* locus and found extensive sRNA mapping confined to the region targeted by the dsRNA vector (Fig. 5A), with the most frequent sRNA length being 20 nt (Fig. 5B-E). Comparison of the TAP treated with untreated samples revealed no difference in the sizes of the mapped RNA sequences (Fig. 5B-E) or their localization across the gene (Fig. 5A). This further supported the hypothesis that there was no canonical RdRP-based amplification of secondary sRNAs.

**Figure 5.**
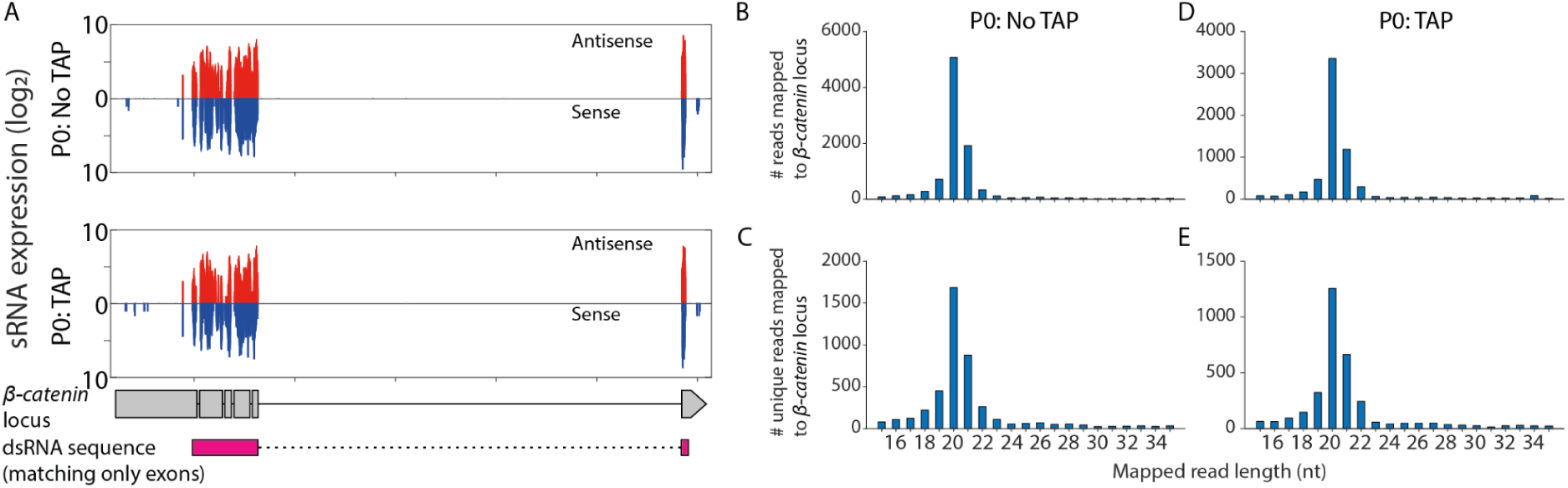
Distribution and characteristics of sRNAs mapping to the *ꞵ-catenin* locus following *ꞵ-catenin* (RNAi). (A) sRNA read distribution across the *ꞵ-catenin* locus in *S. mediterranea*. Normalized read counts for antisense (red) and sense (blue) sRNAs show alignments only at the dsRNA-targeted region. The *ꞵ-catenin* gene structure (exons, gray blocks) and the dsRNA trigger sequence (pink block, only to exonic regions) are mapped below. Libraries were generated either without (top) or with (bottom) TAP treatment. (B–E) Length distribution of sRNAs mapping to the *ꞵ-catenin* locus. A distinct 20–21 nucleotide (nt) peak dominates the sRNA population. This 20–21 nt enrichment is present in both standard (B, C) and TAP-treated (D, E) libraries, and persists whether plotting total mapped reads (B, D) or restricting the analysis to unique read sequences (C, E).

### RNAi memory is characterized by sRNAs containing non-templated 3’ adenosines

So far, we have shown that RNAi-induced silencing is long-lasting (Fig. 1D), post-transcriptional (Fig. 4B), independent of RdRP activity (Fig. 5B-E), and does not require the sustained presence of dsRNA (Fig. 2B). However, the nature of the RNAi-induced molecules that persisted during this prolonged response remained unclear. While we had initial indications that sRNAs were generated, the dynamics and composition of the sRNA pool over time were unknown.

We used dsRNA to silence *ovo-1* and collected sRNA samples for sequencing at three time points (Methods): (i) one week post-feeding (P0); (ii) after one regeneration cycle (F1); and (iii) after three regeneration cycles (F3). We then mapped the resulting sRNA data to the planarian genome, and examined changes in the properties of the sRNAs over time. At the initial time point (P0), the size distribution of sRNAs mapped to the targeted locus was highly varied, with sequences extending up to 47 nucleotides (Fig. 6A, S6A-D) in length and a comparable number of reads aligning to both the sense and antisense strands (Fig. S6A-J). We suspected that many of the reads represented dsRNA degradation products. Indeed, in P0 samples, we found reads corresponding to the terminal adapter sequences present in the dsRNA, which are not part of the planarian genome (Fig. S6K; Methods).

**Figure 6.**
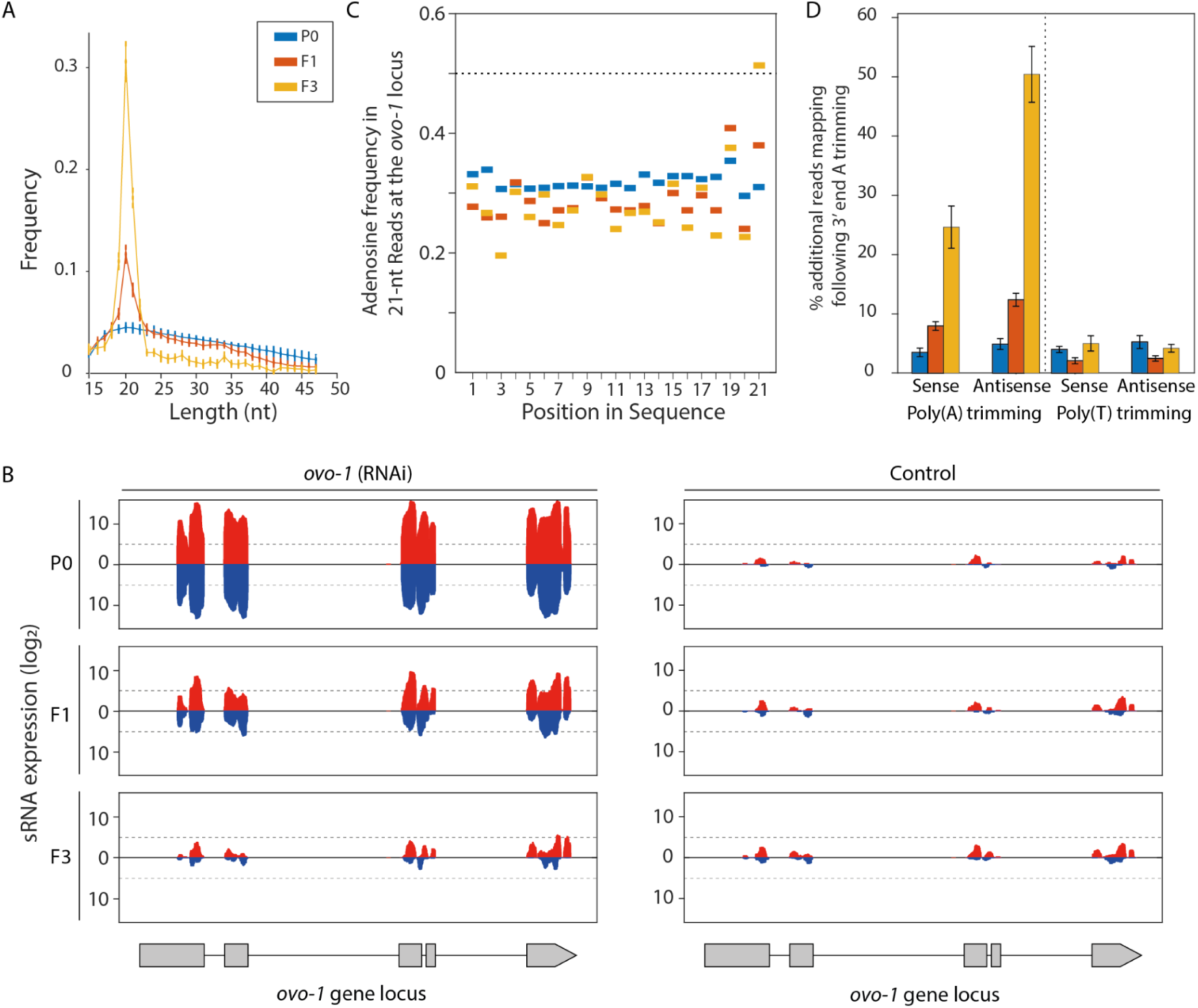
Characteristics of sRNA mapping to the *S. polychroa ovo-1* gene locus following RNAi. (A) sRNA length distribution mapped across the *ovo-1* gene locus at multiple regeneration cycles showing a striking enrichment of 19–22 nt reads in F1 and F3 generations. Data represent the mean of three (P0) or six (F1, F3) biological replicates. Shown are uniquely mapped reads after removing redundant sequences. (B) Small RNA read distribution across the *ovo-1* locus following *ovo-1* (RNAi) (left) or control RNAi (right). Normalized read counts for antisense (red) and sense (blue) sRNAs are plotted against the *ovo-1* gene structure (exons, gray blocks). Data represent the average of three (P0, all controls) or six (F1, F3 *ovo-1* RNAi) replicates. (C) Nucleotide frequency across each position of perfectly mapped 21-nt sRNAs at the *ovo-1* locus. Adenosine frequency at position 21 increases progressively across generations, likely related to the presence of non-templated adenosines at the 3’ end. Reads were pooled across all replicates for each generation. (D) Percentage of additional unique reads (≥15 nt) successfully mapped to the *ovo-1* locus following 3’ poly(A) or poly(T) trimming. The increase in reads requiring poly(A) trimming (left) across regeneration cycles indicates the progressive accumulation of non-templated 3’ A-tails. Poly(T) trimming controls (right) demonstrate this enrichment is specific to A-tailing rather than a general sequence or GC-content bias. Bar height indicates average and error bars the standard deviation.

In the libraries collected following regeneration (i.e., F1 and F3), we observed notable changes in the sRNA characteristics. The total number of reads mapped to the dsRNA-targeted region decreased by roughly 1000-fold (Fig. 6B; S6A-B). Yet, this decline in sRNA abundance was not uniform across all sRNA sizes (Fig. 6A; S6A-J). The relative abundance of 19–22 nt sRNAs mapped to the *ovo-1* locus following *ovo-1* (RNAi) significantly increased between the initial and later time points, with longer sequences largely depleted from these samples (Fig. 6A; S6A-D). Moreover, we observed a consistent enrichment of sRNAs mapped to the antisense strand compared to the sense strand (Fig. S6I).

Notably, the sRNAs targeting *ovo-1* carried a non-templated ’A-tail’ of one to three adenosine residues at their 3ʹ end, which limited read mapping prior to 3’ poly(A) trimming (Fig. 6C-D). This 3’-end A enrichment was absent in the libraries collected at the initial (P0) time point (Fig. 6D), indicating that it was added to sRNAs over generations. By contrast, sRNAs mapping to untargeted genomic loci lacked this modification (Fig. S6L). This specificity suggested that exposure to exogenous dsRNA triggered a mechanism that selectively adenylates target-directed sRNAs. Because this non-templated 3’ adenylation accumulated over generations, it raised the possibility that such processing is associated with the long-term persistence of RNAi.

### Novel RNAi sensor reveals that long-term RNAi memory requires an endogenously encoded target

There are currently no genetically encoded reporters for gene activity in planarians. To explore the characteristics of RNAi memory, we developed a novel RNAi sensor by taking advantage of the recently described RNA-based nanoluciferase (Nluc) expression assay ^34,35^ (Fig. 7A). Specifically, our sensors comprised the Nluc open reading frame (ORF), flanked by specific 3’UTR sequences as targets of exogenously supplied dsRNA (Fig. 7A).

**Figure 7.**
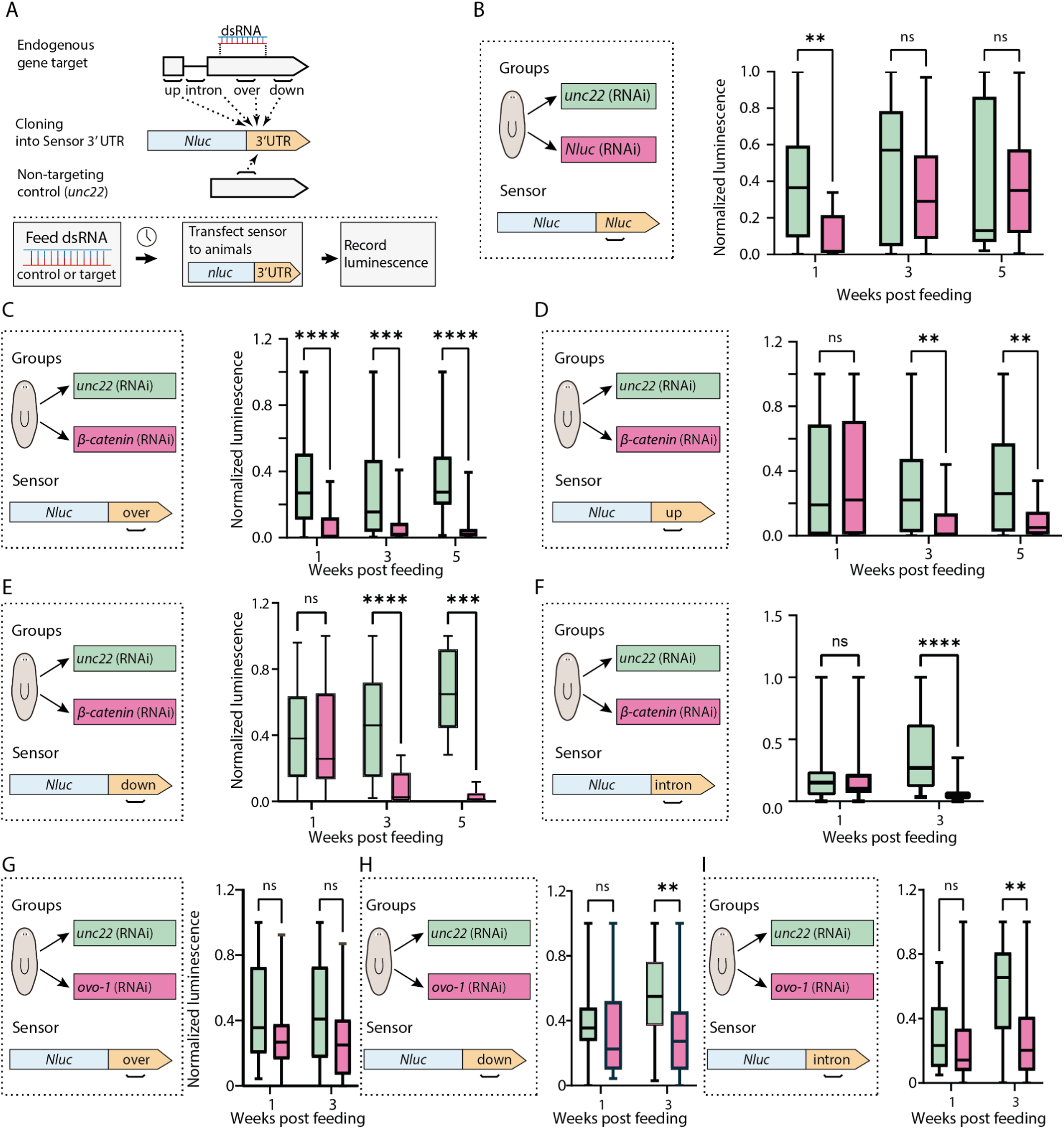
RNAi-mediated gene silencing acts in *trans* and spreads across the target locus. (A) Schematic of the *in vivo* Nluc sensor assay. Animals are fed dsRNA targeting an endogenous gene or a non-targeting control (*unc22*), then transfected with an *Nluc* sensor mRNA. The sensor 3ʹ UTR contains target gene sequences located upstream (up), overlapping (over), downstream (down), or intronic to the exogenous dsRNA trigger. (B) Normalized luminescence of the Nluc sensor following *Nluc* or control (RNAi). Exogenous *Nluc* dsRNA robustly suppressed the sensor when transfected 1 week post-feeding, but this suppression is lost by weeks 3 and 5. (C–F) Trans-silencing dynamics across the *ꞵ-catenin* locus. Nluc sensors containing *ꞵ-catenin* sequences that (C) overlap the dsRNA target maintain significant suppression at 1, 3, and 5 weeks. By contrast, sensors carrying (D) upstream, (E) downstream, or (F) intronic sequences escape silencing at 1 week, but undergo significant suppression at weeks 3 and 5 (intronic measured only at weeks 1 and 3). (G–I)Trans-silencing dynamics across the *ovo-1* locus following a brief, two-feed RNAi course. The (G) overlapping sensor shows no significant reduction in luminescence. Conversely, sensors containing (H) downstream and (I) intronic sequences exhibit significant suppression at 3 weeks post-feeding, demonstrating that silencing spreads to untargeted regions even when the primary target region escapes sustained suppression. Data are represented as box plots where whiskers indicate minimum and maximum values, horizontal bar the median, and the box the interquartile range (IQR). For all panels, 8–12 animals were injected per group per experiment. Experiments were repeated at least twice, yielding 3–4 biological replicates for most groups. Statistical significance was calculated using multiple Mann-Whitney U tests with Holm-Sidak correction (ns = not significant; *P < 0.05; **P < 0.01; ***P < 0.001; ****P < 0.0001).

To investigate the temporal dynamics of silencing, we fed animals *Nluc* or control (*unc-22*) dsRNA and subsequently injected *Nluc* reporter mRNA at 1, 3, or 5 weeks post-feeding. At one week, *Nluc* dsRNA treatment robustly suppressed luminescence relative to controls, confirming the efficacy of the initial RNAi response (Fig. 7B). By contrast, there was no significant reduction in luminescence at 3 or 5 weeks post-RNAi (Fig. 7B).

Because RNAi targeting endogenous genes produced sustained silencing (Fig. 4B), we next investigated whether dsRNA directed against such targets produced longer Nluc suppression. We fed animals *ꞵ-catenin* or control dsRNA. After confirming that *ꞵ-catenin* RNAi did not generally compromise the Nluc reporter (Fig. S7A), we injected chimeric *Nluc* mRNAs containing *ꞵ-catenin* 3’UTRs at 1, 3, and 5 weeks (Fig. 7A, C). The *Nluc*-*ꞵ*^over^ construct (Fig. 7C), which overlaps the dsRNA sequence, showed significant suppression at all time points (Fig. 7C). This effect was strictly sequence-specific: *ꞵ-catenin* (RNAi) animals did not suppress *Nluc* transcripts containing an *ovo-1* 3’UTR (Fig. S7B). These data indicate that RNAi memory involves sustained, sequence-specific suppression against endogenous targets (Fig. 7C). While abundant dsRNA likely drives the initial silencing (Phase 1), suppression at 3 and 5 weeks (Phase 2) persists only for the endogenous sequence (Fig. 7B-C). This specific persistence likely marks the transition from direct dsRNA-mediated silencing to a stable memory state.

### RNAi-induced *trans-*silencing is guided by information spanning the target gene locus

So far, we showed that RNAi produces a sustained decrease in gene expression that was accompanied by antisense sRNAs and in particular sRNAs with non-templated ’A-tails’ at their 3ʹ end. Results from the Nluc RNAi sensor (Fig. 7B-C) indicated the existence of a *trans*-acting molecule that maintains silencing after the exogenous dsRNA has disappeared. To identify the properties of this heritable agent, we initially suppressed three genes: *ꞵ-catenin*, *ovo-1*, and a non-targeting control (*unc-22*) (Methods), and examined the suppression of a panel of Nluc sensors whose 3’ UTRs contained different sequences (Fig. 7C-I).

We did not observe the production of sRNAs outside the dsRNA-targeted region (Fig. 5A, 6B). Therefore, we predicted that Nluc sensors containing *ꞵ-catenin* sequences outside this targeted region would not be silenced. We engineered Nluc reporters carrying *ꞵ-catenin* sequences that either overlapped (ꞵ^over^), or were located upstream (ꞵ^up^) or downstream (ꞵ^down^) of the exogenously supplied dsRNA trigger sequence (Fig. S7C). We fed animals *ꞵ-catenin* dsRNA and assayed them at 1, 3, and 5 weeks. At each time point, we transfected animals with one of the *Nluc* constructs and measured luminescence (Fig. 7A; Methods).

As described above, the reporter containing the overlapping 3’ UTR (ꞵ^over^) was suppressed at all time points (Fig. 7C). By contrast, at one week post-feeding, neither the ꞵ^up^ nor the ꞵ^down^ sensor was silenced, indicating that the initial dsRNA trigger did not target adjacent regions of the transcript (Fig. 7D-E). Strikingly, this changed at later time points: by week 3, both ꞵ^up^ and ꞵ^down^ were repressed as strongly as ꞵ^over^, and this repression persisted in week 5 (Fig. 7D-E). Thus, between weeks 1 and 3, cells acquired the capacity to recognize *ꞵ-catenin* sequences outside the original dsRNA trigger, demonstrating a spread of the silencing effect. A control reporter containing an *ovo-1* 3ʹ UTR remained active at all time points, confirming target locus specificity (Fig. S7B).

Next, we designed an *Nluc* construct (ꞵ^intron^) that included an intronic sequence derived from the *ꞵ-catenin* locus in the 3’ UTR (Fig. S7C). We transfected the construct into *ꞵ-catenin* (RNAi) animals at two time points (1 and 3 weeks). At the initial time point, we did not observe silencing of the Nluc sensor (Fig 7F). However, as we observed upon the inclusion of exonic sequences, we detected strong suppression of the reporter at the later time point (3 weeks; Fig. 7F). Importantly, dsRNA targeting the *ꞵ-catenin* intron directly did not induce gene silencing (Fig. S7D), suggesting that propagation is unidirectional (i.e., exon to intron). Together, the data suggested that the *trans*-acting silencing signal triggered by RNAi recognized any sequence derived from the *ꞵ-catenin* locus, including regions absent from the mature mRNA.

To generalize our findings, we analyzed trans-silencing after targeting a different gene, *ovo-1*. We prepared three Nluc reporters containing 3’ UTRs that either overlapped the *ovo-1* dsRNA sequence (o^over^), localized to a downstream exon (o^down^), or localized to an intron (o^intron^). We fed animals *ovo-1* dsRNA twice, a short RNAi course that does not produce a loss-of-eye phenotype. We then transfected the *ovo-1* (RNAi) animals with the Nluc reporters. At 1 week post-RNAi, we observed no suppression of any construct, consistent with the lack of visible RNAi effects. At week 3, the o^over^ reporter remained active (Fig. 7G), but the o^down^ reporter exhibited significant suppression (Fig. 7H). Furthermore, the o^intron^ reporter showed strong silencing at week 3, indicating that the silencing effect extended from the initially targeted region to intronic sequences (Fig. 7I). We observed similar intronic reporter silencing when targeting *caspase-8* (Fig. S7F-G). These results demonstrate a dynamic RNAi response comprising distinct phases: a priming phase dependent on the dsRNA or its direct products, and a subsequent, sustained phase characterized by broader target recognition across the gene locus in the absence of the original dsRNA.

## Discussion

Our results demonstrate that planarians sustain potent, sequence-specific RNA interference for at least three months and across multiple cycles of whole-body regeneration, a remarkable capacity in an organism whose tissues are continually renewed by pluripotent stem cells ^1,2^. To explain this long-term persistence, we propose a two-phase model of RNAi memory. In phase 1 (the priming phase), which occurs immediately following dsRNA ingestion, dsRNA targeting either endogenous (e.g., *ꞵ*-catenin) or exogenous (e.g., *Nluc*) sequences efficiently suppresses gene expression. This phase is characterized by the relative abundance of exogenous dsRNA (as observed via qPCR) and the horizontal transferability of RNAi-mediating factors (as evidenced by our 1-week transplantation assays). During this phase, silencing of exogenous sequences is likely achieved through sRNAs that were directly derived from the ingested dsRNA, and the only requirement for silencing at this phase is sequence complementarity between the target and dsRNA. RNAi spreading has been recently proposed to happen via extracellular vesicles ^24^. However, the priming phase is limited in duration by the short dsRNA half-life.

During phase 2 (the memory phase), silencing spreads bidirectionally from the originally targeted region into adjacent, non-targeted regions of the gene sequences. At this stage, dsRNA is largely absent, and RNAi memory is maintained cell autonomously. Importantly, this persistence requires that the dsRNA target an endogenously expressed gene. Since silencing propagates outside the targeted region, it is likely driven by endogenously produced molecules, although our sRNA sequencing analyses could not identify their precise identity. We propose that the potent, long-lasting RNAi observed in planarians reflects an early contribution from dsRNA-derived sRNAs that is later reinforced by endogenous factors. Given that planarians lack canonical RdRPs and the spreading encompasses both exons and introns, we infer that RNAi memory is maintained through a previously unrecognized, RdRP-independent mechanism.

In *C. elegans*, transgenerational silencing depends on RdRP-derived 22G sRNAs and nuclear Argonaute proteins ^36,37^. Our analyses showed that planarians, by contrast, lack identifiable eukaryotic RdRP genes, display no 5ʹ-triphosphorylated sRNAs, and show no change in chromatin accessibility or pre-mRNA output at the targeted locus. Instead, silencing is maintained post-transcriptionally: mature mRNA is depleted while pre-mRNA levels remain stable. Moreover, our *Nluc* sensor system confirmed that this repression likely operates exclusively in the cytoplasm. These features place planarians among the few metazoans capable of persistent RNAi, stable in the parent and its asexual progeny, without RdRPs, suggesting that the ancestral bilaterian toolkit included multiple, mechanistically distinct strategies for sRNA-based memory of RNAi.

Three observations further refine this model. First, after an initial dependence on exogenous dsRNA for suppression of gene expression, the silencing signal spreads from the originally targeted exonic sequences to other regions in the locus, including exons and introns. Second, this spreading coincides with the appearance of ∼20 nt antisense sRNAs, modified by non-templated 3ʹ adenosine tails. Such 3’ end tails are recognized in other systems for altering the stability of certain sRNAs, particularly miRNAs, either positively ^38^ or negatively ^39^. Third, there is a brief, one-week window after dsRNA exposure during which the silencing signal can transfer horizontally from host tissue to a naive graft; afterwards, it is inherited vertically in a cell-autonomous manner. These findings explain why our RNAi sensor detected durable repression only when its reporter contained an endogenous sequence: an exogenous target is silenced transiently by the initial dsRNA-derived effectors, whereas an endogenous target can further supply substrates that generate unknown secondary silencing molecules necessary for long-term maintenance. We therefore propose a two-phase process: an early, mobile population of primary effectors initiates silencing throughout the animal, followed by a stable, self-sustaining pool of silencing effectors derived from the endogenous locus that preserves repression in descendant cells.

The existence of such an RdRP-independent memory system raises questions about its physiological role. One likely function, conserved across RNA-based pathways, is defense against selfish genetic elements such as transposons and viruses. The spreading across the endogenous template could also be used to further suppress sequences proximal to the selfish element. However, this requirement for an endogenous template, could also suggest that the pathway has been co-opted for developmental gene regulation. By establishing a long-lasting post-transcriptional record of a gene activity state, this mechanism could stabilize cell fates and ensure correct tissue patterning during the continual cell turnover that accompanies growth, homeostasis, and regeneration. It appears finely tuned to discriminate self from non-self: any sequence can elicit a transient response, but only transcripts produced within the worm can induce long-term memory, thereby limiting the duration of silencing triggered by environmental RNA.

Are these sRNAs amplified in the absence of RdRP? Our study does not provide an answer to this question, but one possibility is that a planarian-specific nucleotidyl-transferase adds protective A-tails we detect. Indeed, planarian genomes encode several non-canonical poly(A) polymerases, which in mammals ^38^ and flies ^39^ have been shown to modify the 3’ ends of miRNAs. Another possibility is that certain PIWI- or AGO-clade proteins in flatworms engage in an RNA-templated cleavage cycle that eliminates the need for a dedicated polymerase ^33^. The eventual extension of repression into intronic regions, which are absent from the mature mRNA, implies an interaction with nascent pre-mRNA in the nucleus. Biochemical isolation of the tailing activity and genetic disruption of candidate factors will be essential to test these ideas.

Our work also intersects with the long-standing, though controversial, literature on memory transfer in planarians. McConnell’s 1960s claim that learned behavior could be conveyed by RNA lacked a molecular substrate. Although we do not address behavioral memory, we show that planarian tissues package gene-specific information into RNA species that persist through drastic cellular turnover and regeneration, making a molecular link between RNA and memory a now-testable hypothesis.

Practically, exposure to dsRNA is sufficient to achieve months-long knockdown, informing functional genomic studies in flatworms and potentially indicating RNA-based control strategies against parasitic relatives. Important questions remain: we have yet to identify the effector molecules that facilitate this long-term silencing. Cell-type-specific sRNA profiling and proteomic capture of Argonaute-bound RNAs will aid in resolving these gaps.

In summary, planarians possess a durable, locus-wide RNAi memory that operates without RdRPs or chromatin modifications and relies on long-lived antisense sRNAs. This work broadens the repertoire of epigenetic inheritance mechanisms in animals and establishes planarians as a powerful model for uncovering how RNA alone can encode cellular memory amid continuous somatic renewal.

## Materials and Methods

### Animal Husbandry

Experiments were performed using clonal strains of asexual *S. mediterranea* and sexual *S. polychroa*. Standard culture conditions, prior to RNAi experiments, maintained the animals at 20 °C in the dark with weekly beef liver feedings. All gene inhibition protocols required a 7-day starvation period immediately prior to experimental onset.

### TA cloning of plasmids for dsRNA synthesis

Planarian cDNA was generated from total RNA using the RevertAid First Strand cDNA Synthesis Kit (Thermo Scientific™; CAT# K1621). Target genes were amplified by PCR using gene-specific primers, and the resulting amplicons were cloned into the pGEM-T vector according to the manufacturer’s instructions (Promega; CAT# A1360). Primer sequences are found in Table S2.

### Bacterial transformation with plasmids

Recombinant plasmids were introduced into *E. coli* TOP10 cells (Thermo Fisher Scientific) via heat-shock transformation. Briefly, 100 μl of competent cells were combined with 5 μl of each ligation reaction, incubated on ice for 30 min, and heat-shocked at 42 °C for 45 seconds. Transformed cells were recovered in 350 μl of LB medium at 37 °C for 1 hour, then plated on agar plates containing ampicillin, isopropyl ꞵ-D-1-thiogalactopyranoside (IPTG), and 5-bromo-4-chloro-3-indolyl-ꞵ-D-galactopyranoside (X-gal). Plates were incubated overnight at 37 °C. Purified plasmids were sequence-verified by Sanger sequencing or Illumina-based sequencing using Plasmidsaurus.

### Cloning of *Nluc* constructs by HiFi DNA Assembly

Various sequences were cloned into a plasmid obtained from the Rink laboratory. The plasmid contains a T7 promoter upstream of a NanoLuc (*Nluc*) reporter gene flanked by 5ʹ and 3ʹ untranslated regions (UTRs), as well as a kanamycin resistance cassette for selection. The vector backbone and insert were linearized by PCR using overlapping primers designed in SnapGene. DNA fragments were assembled using NEBuilder® HiFi DNA Assembly (NEB; CAT# E2621L) according to the manufacturer’s instructions. Briefly, PCR-amplified vector and insert concentrations were used to calculate molar ratios, and fragments were combined at a 1:2 vector-to-insert molar ratio. 10 µL of NEBuilder HiFi DNA Assembly Master Mix was added, with the reaction volume being adjusted to 20 µL with nuclease-free water. The assembly reaction was incubated at 50 °C for 15 minutes prior to bacterial transformation, as described above.

### Double-stranded RNA synthesis for RNAi experiments

Double-stranded RNA (dsRNA) was synthesized as previously described ^6^. Briefly, DNA templates for *in vitro* transcription (IVT) were generated through PCR amplification of cloned sequences using primers that included 5’ flanking T7 promoter sequences. dsRNA was synthesized using either the TranscriptAid T7 High Yield Transcription Kit (CAT# K0441; Thermo Scientific) or HiScribe T7 High Yield RNA Synthesis Kit (NEB; CAT# E2040S). IVT reactions were incubated overnight at 37 °C and then treated with RNase-free DNase (supplied in the kit) for 30 minutes. dsRNA was purified by ethanol precipitation and finally resuspended in 60 µl of ultrapure water. RNA was run on 1% agarose gels and quantified by Qubit 4, using the RNA broad-range kit. Animals were starved for a week before starting RNAi experiments. *Schmidtea mediterranea* (Asexual) or *Schmidtea polychroa* (Sexual) were used in all experiments.

Animals were fed dsRNA mixed with beef liver twice weekly, as previously described ^40^. For *ovo-1* (RNAi), *S. polychroa* were fed *ovo-1* dsRNA 12 times. For *ꞵ-catenin* (RNAi), *S. mediterranea* were fed *ꞵ-catenin* dsRNA twice. As a control, animals were fed *unc-22* dsRNA, a gene that is absent in planarians. *S. mediterranea* were used to perform the *Nluc* RNAi-sensor experiments. For these experiments, *Nluc* (RNAi), *ꞵ-catenin* (RNAi), *ovo-1* (RNAi) and *caspase-8* (RNAi) animals were fed twice with the corresponding dsRNA.

### Fluorescence-activated cell sorting (FACS)

Planarians were cut to fragments using a blade. Tissue fragments were collected into calcium-free, magnesium-free medium plus 0.1% BSA (CMFB) and dissociated by pipetting vigorously for 5-10 minutes. To reduce clumping at downstream steps, dissociated cells were strained through a 40 μm filter. Cells were centrifuged at 315 G for 5 minutes at 4 °C and resuspended in CMFB containing Hoechst 33342 (40 μl/ml) for 45 min at RT in the dark. Before loading into the FACSAria II flow cytometer (Becton Dickinson), cells were labeled with propidium iodide (5 μg/ml) for cell viability detection. FACS gating was performed as previously described for planarian cell populations ^41^. Cells were sorted into Trizol LS (Sigma-Aldrich; CAT# T3934-100ML).

### RNA extraction

Animals were homogenized in 700 μl of TRI Reagent (Sigma; CAT# 9424) using 0.5 mm zirconium beads in a bead-beating homogenizer (Allsheng; Bioprep-24) for two 45-second cycles, with a 20-second interval between cycles. Following homogenization, samples were incubated at room temperature for 5 minutes. Chloroform (140 μl) was then added to each tube, which was shaken vigorously for 15 seconds and incubated at room temperature for an additional 3 minutes. Samples were centrifuged at 12,000 G for 25 minutes at 4 °C, after which the aqueous phase was transferred to a new tube. Isopropanol (500 μl) was added, and tubes were inverted five times and incubated at room temperature for 10 min. Samples were centrifuged at 12,000 G for 45 minutes at 4 °C. The supernatant was discarded, and the RNA pellet was washed twice with 75% ethanol, followed by centrifugation at 7,500 G for 5 minutes at 4 °C. Pellets were air-dried for 10 minutes and resuspended in 32 μl of nuclease-free water. RNA concentration was quantified using a Qubit fluorometer (Invitrogen; Q33226) according to the manufacturer’s instructions.

### cDNA synthesis from mRNA and total RNA

We synthesized cDNA from RNA using the RevertAid H Minus First Strand cDNA Synthesis Kit as per the manufacturer’s protocol (Thermo Scientific; CAT# K1631). Random hexamer primers were used to generate cDNA from total RNA, whereas oligo(dT) primers were used for adenylated RNA (mRNA). The resulting cDNA was subsequently used as a template for qPCR.

### RT-qPCR and analysis

qPCR was performed with 2x qPCRBIO Fast qPCR SyGreen Blue Mix (PCRBIO; CAT# PB20.15-20). The expression of the target genes was measured using the QuantStudio 3 Real-Time PCR system (Applied Biosystems) using the following program [95 °C for 20 seconds, 40 cycles (95 °C for 1 second, 60 °C for 20 seconds)], with two technical replicates per sample and at least two biological replicates per sample. The relative gene expression fold-change was calculated by the ΔΔCt method with *gapdh* used as an endogenous control. Forward and reverse primers were designed for each target gene. Primer efficiency was tested prior to the experiment by using the standard curve method for five decreasing cDNA concentrations.

### qPCR to detect total dsRNA and full-length dsRNA

To detect dsRNA species, qPCR assays were designed to distinguish between total and full-length dsRNA. For total dsRNA detection, cDNA was synthesized using random hexamers, and qPCR primers were designed to anneal to a dsRNA-specific adapter sequence and a downstream region. Because these adapter sequences are absent from planarian genomic and transcriptomic sequences, amplification occurs only in the presence of dsRNA. To specifically detect full-length dsRNA, cDNA was synthesized using a gene-specific reverse transcription primer that anneals to one adapter, followed by qPCR using primers targeting the second adapter and a downstream region. Amplification is therefore restricted to cDNA derived from intact, full-length dsRNA molecules.

### qPCR to detect mature mRNA and pre-mRNA

Mature mRNA and pre-mRNA were distinguished by qPCR primer design based on transcript splicing. Mature mRNA was detected using exon–exon spanning primers positioned outside the dsRNA target region, whereas pre-mRNA was detected using intron-specific primers. Genomic DNA contamination was minimal, as no-RT controls exhibited Ct values approximately 10 cycles higher than reverse-transcribed samples. Detection of pre-mRNA was performed following cDNA synthesis using random hexamers.

### Transplantation and tissue grafting

Grafting was performed as previously described ^42^, with minor modifications. Animals were anesthetized in 0.1% (w/v) chloretone for 3–4 minutes, rinsed in 1× modified Holtfreter’s solution (MHS), and placed in a chilled Petri dish lined with Grade 2 Whatman filter paper soaked in MHS. Tissue plugs from regions below the eyes and above the pharynx were excised from donor animals using glass capillaries with a larger internal diameter (ID 0.80 × 0.80 mm; CM Scientific; CAT# 8280-50) and positioned onto host animals using forceps. Corresponding tissue below the eyes was removed from host animals using a smaller internal diameter glass capillary (ID 0.50 × 0.50 mm; CM Scientific; CAT# 8250-50) and discarded, after which donor tissue plugs were inserted into the cavity. Grafted animals were transferred to smaller Petri dishes lined with MHS-soaked Grade 2 Whatman filter paper, covered with moist cigarette paper, and incubated at 10 °C for 12 hours. Following incubation, the cigarette paper was removed and animals were transferred to planarian water.

### Lethal exposure to γ-Irradiation

Animals were lethally irradiated using a BIOBEAM GM 8000 (Gamma-Service Medical GmbH) with 6000 rads ^43,44^. Grafting experiments were performed 24 hours following irradiation.

### ATAC-seq and analysis

ATAC-seq analysis was done for *ꞵ-catenin* and *unc-22* RNAi *S. mediterranea* worms as previously described ^27^. To summarize, planarian mucus was first stripped using 0.5% (w/v) N-acetyl-L-cysteine (CAS: 616-91-1) in 20 mM HEPES-NaOH (pH 7.25), followed by tissue pulverization in liquid nitrogen. Samples were stored at −80 °C. Nuclei were isolated in a chilled homogenization buffer [260 mM Sucrose, 30 mM KCl, 10 mM MgCl₂, 20 mM Tris-HCl (pH 8)] and purified using a 60% iodixanol (OptiPrep, Sigma-Aldrich, D1556-250ML) density gradient. After mechanical dissociation with a Dounce homogenizer (pestle B, Cat# D8938-1SET) and filtration through 50 µm CellTrics filters (Cat# 04-004-2327), approximately 50,000 nuclei per sample were tagmented at 37 °C for 30 minutes. The reaction utilized the Illumina Tagment DNA Enzyme and Buffer Large Kit (Cat# 20034198), supplemented with 1% Digitonin, 10% Tween-20, and 53.5 mM DTT. Tagmented DNA was purified using the Zymo DNA Clean & Concentrator-5 kit (Cat# D4014) and PCR-amplified to generate sequencing libraries, which were validated by assessing fragment size distribution using an Agilent 2100 Bioanalyzer.

Following sequencing, ATAC-seq reads were first trimmed using Cutadapt ^45^, and quality control was performed with FastQC ^46^ prior to mapping. Trimmed reads were aligned to the *S. mediterranea* genome (schMedS1-sexual-GO-v1 assembly) ^30^ using BWA (v0.7.17; ^47^). PCR duplicates were removed with Picard (v2.21.4). Peak calling and ATAC signal quantification were performed using MACS2 (v2.2.9.1; ^48^). Significant peaks (FDR < 0.05) were filtered and merged within each sample using Bedtools. Candidate peaks were then compiled across all samples, and sample support was quantified using Bedtools intersect with a ≥40% reciprocal overlap threshold. Peaks supported by all 18 samples were retained as consensus peaks. The number of merged peaks per sample ranged from 64,500-175,297, resulting in 19,428 consensus peaks.

### sRNA sequencing and analysis

The NEBNext Multiplex Small RNA Library Prep Set for Illumina from New England Biolabs was used for small RNA library preparation, following the manufacturer’s protocol. The concentration of the libraries was determined using Qubit, and their quality was assessed using an Agilent 2200 TapeStation. The libraries were pooled and electrophoresed on a 4% agarose E-Gel from Life Technologies. Fragments ranging in length from 140 to 160 nt were carefully excised and purified using the MinElute Gel Extraction Kit (QIAGEN). The purified pool was assessed for concentration and quality using Qubit and the Agilent 2200 TapeStation. Libraries were sequenced on an Illumina NextSeq 500 sequencer at the CRCF at the Faculty of Life Sciences in Tel Aviv University. Bioinformatic analysis was performed by assessing the resultant fastq quality using FastQC ^46^. The sequencing reads were then subjected to adapter clipping using Cutadapt ^45^. Next, the clipped reads were mapped to the schMedS1-sexual-GO-v1 and the schPol2 assembly versions of *S. mediterranea* and *S. polychroa* genomes, respectively, using ShortStack ^49^. Gene-level read counts were obtained using the Python-based script HTSeq-count ^50^.

### Computational analysis of RdRP domain in planarian transcriptomes

Sequences predicted to encode the eukaryotic-type RdRP domain (PF05183) in *M. lignano* were identified using InterPro ^51^. A total of 23 potentially redundant protein sequences were retrieved (IDs: A0A1I8FC89, A0A1I8FH82, A0A1I8FNC6, A0A1I8GGK4, A0A1I8GH91, A0A1I8H5L0, A0A1I8H6D6, A0A1I8HJL9, A0A1I8IKM0, A0A1I8ILF0, A0A1I8INY5, A0A1I8IPU0, A0A1I8IQ63, A0A1I8JJ98, A0A1I8JQW4, A0A267EGF3, A0A267FD57, A0A267FEP9, A0A267FM32, A0A267FVJ0, A0A267G4F5, A0A267GS89, A0A267H6U2). The sequences were queried against predicted translation of planarian transcriptome assemblies from the PlanMine repository ^30^ using TBLASTN (v 2.17.0+) ^52^. Resulting hits with an E-value < 10^-5^ were scanned for the PF05183 domain. This domain was not detected in any of the hits; these sequences likely represent other regions that bind or operate on RNA. Additionally, planarian transcriptome assemblies ^30,53^ were searched directly for RdRP-encoding domains. Homology searches were performed using BLAST+ (v 2.17.0+) against the PlanMine transcriptomes using known RdRP protein sequences: EGO-1 from *C. elegans* (G5EBQ3), RDP1 from *Schizosaccharomyces pombe* (O14227), and RDR6 from *Arabidopsis thaliana* (Q9SG02). No significant hits were identified. Finally, available transcriptome-wide Pfam annotations ^30,53^ were examined for eukaryotic RdRP protein domains, yielding no significant hits.

### Synthesizing 5’ capped and 3’ A-tailed mRNA for transfection

Capped and A-tailed mRNA was synthesized as previously described ^34,35^. Briefly, the sequence to be synthesized was amplified by PCR. Reaction cleanup was performed using Wizard® SV Gel and PCR

Clean-up System (Promega; CAT# A9282). The amplified DNA included a T7 promoter. Then, using T7 mScript™ Standard mRNA Production System V2 (CellScript; CAT# C-MSC11610), RNA was synthesized for 30 minutes at 37 °C. The samples were then treated with included DNase for 15 minutes at 37 °C. Phenol-chloroform extraction is performed on all samples. 5’ capping was done with the ScriptCap Capping Enzyme and ScriptCap 2’-O-Methyltransferase for 30 minutes at 37 °C. Next, 3’-poly(A) tailing was performed on the RNA samples for 30 minutes at 37 °C. Next, the RNA samples were purified by phenol-chloroform extraction and the concentration was quantified using a Qubit fluorometer (Invitrogen; Q33226). The mRNA was diluted to a concentration 0.5 µg/µl with nuclease-free water, aliquoted and stored at −80 °C.

### Transfection of *Nluc* sensors

mRNA transfection was performed as previously described ^34,35^ using the TransIT®-mRNA Transfection Kit (Mirus Bio; CAT# 2225). Briefly, 2 µL of TransIT-mRNA transfection reagent, 1 µL of TransIT Boost reagent, and 3 µL (1.5 µg) of mRNA were combined in a PCR tube and brought to a final volume of 25 µL with serum-free L-15 medium (Thermo Scientific; CAT# 11415049). The mixture was incubated for 5 minutes at room temperature. Subsequently, 8 µL of the mixture was loaded into a pulled glass needle using Femtotips Microloader Tips (Eppendorf; CAT# E5242956003). Worms were positioned ventral side up on black filter paper and chilled on a Peltier cooling plate. Fine forceps broke the needle tip to establish the injection aperture. A separate stabbing needle pre-wounded each animal between the tail tip and the posterior end of the pharynx. A micromanipulator guided the transfection needle into the pre-existing wound, and a FemtoJet microinjector delivered approximately 0.3 µL of the mixture into each animal.

### Detection and analysis of luminescence

Detection of luminescence was done as previously described ^34,35^. In summary, injected worms were finely macerated with a razor blade. The tissue was suspended in 250 µl of L-15 and pipetted into a white opaque flat-bottom 96 well plate (Greiner Bio-one; CAT# 655083). For each reaction (1 reaction per worm), the Nano-Glo Lysis mix was prepared by combining 2 μL of Nano-Glo Luciferase Assay Substrate with 98 μL of Nano-Glo Luciferase Assay Buffer (Promega; CAT# N1130). Immediately after preparation, 100 μL of the lysis mix was added to each well containing dissociated tissue. The mix was gently pipetted up and down 10 times to ensure thorough mixing. For efficiency, a multi-channel pipettor was used. Luminescence was detected on a plate reader (BioTek Synergy 2) immediately after adding the lysis mix. The luminescence values were converted to scores that range from 0 to 1 using the simple formula below:

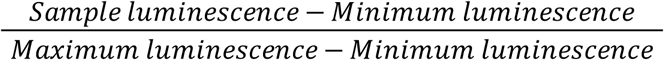

This was performed independently for each *Nluc* construct. Hypothesis testing was performed using two-tailed Mann-Whitney U tests, as the data did not follow a normal distribution.

### Imaging

Fluorescence and confocal images were acquired using a Zeiss LSM800 confocal microscope. Live images were captured with a Leica S9i stereomicroscope. Images were processed in Adobe Photoshop, defining planarian outlines with the lasso and object selection tools. Image analysis was performed using Fiji ^54^.

### DIG-labeled RNAi and immunofluorescence

Digoxigenin (DIG)-labeled *β-catenin* dsRNA was synthesized using DIG-11-UTP nucleotides (Sigma-Aldrich; CAT# 3359247910). To control for labeling specificity, matching control animals were fed unlabeled *β-catenin* dsRNA. For phenotypic observation, a subset of animals was amputated 24 hours prior to dsRNA feeding and assessed for the *β-catenin* (RNAi) phenotype. The remaining animals were fixed at 3, 7, and 14 days post-feeding as previously described ^55^. Briefly, animals were killed using a 5% N-acetyl-L-cysteine (NAC; Mercury, CAT# 1124220100) solution in PBS for 5 minutes. Tissues were then incubated in 4% formaldehyde (FA) in 0.3% PBSTx (phosphate-buffered saline, 0.3% Triton X-100) for 20 minutes. Following fixation, the animals were washed in PBSTx and a 50:50 PBSTx:methanol solution, and stored in 100% methanol at −20 °C.

For immunofluorescence (IF), fixed samples were bleached as previously described ^55^ and permeabilized with proteinase K (2 μg/mL; Invitrogen, CAT# 25530-049) in 0.3% PBSTx. Samples were blocked for 2 hours in 0.5% Roche Western Blocking Reagent (Sigma-Aldrich; CAT# 11921673001) and 5% heat-inactivated horse serum (Biological Industries; CAT# 04-124-1A) diluted in TNTx (100 mM Tris pH 7.5, 150 mM NaCl, 0.3% Triton X-100). The animals were then incubated with an anti-DIG-POD antibody (1:1,500; Roche, CAT# 11207733910) overnight at 4 °C. Following PBSTx washes, tyramide signal amplification was performed as previously described ^55^. Finally, the samples were counterstained with DAPI (1:5,000 in PBSTx) overnight at 4 °C and mounted using Vectashield (Vector Laboratories; CAT# H-1000-10).

### Data availability

Sequencing data produced here, including sRNA libraries and ATAC-seq, is to be released via the Sequence Read Archive (SRA) ^56^ with BioProject accession PRJNA1435607.

## Acknowledgments

We thank all the Wurtzel, Rechavi, and Rink lab members for their helpful comments and critical input. We thank Fruzsina Ficze for the preparation of ATAC-seq libraries. The work was supported by Volkswagen Foundation / Land Niedersachen research cooperation with Israel, SPRUNG (no. 76251-4657/2022, ZN 4035). O.W. is supported by the Israel Science Foundation (no. 2741/25) and the European Research Council (no. 853640). O.W. is a Zuckerman Faculty Scholar. O.R. is grateful for funding from the European Research Council (no. 819151), the Eric and Wendy Schmidt Fund for Strategic Innovation (Schmidt Sciences Polymath Award no. 0140001000), the Khan Foundation (grant no. 0604918421) and the Deutsche Forschungsgemeinschaft (no. 0604918111).

## Competing interests

The authors declare no competing interests.

## Supplementary Figures

**Figure S1.**
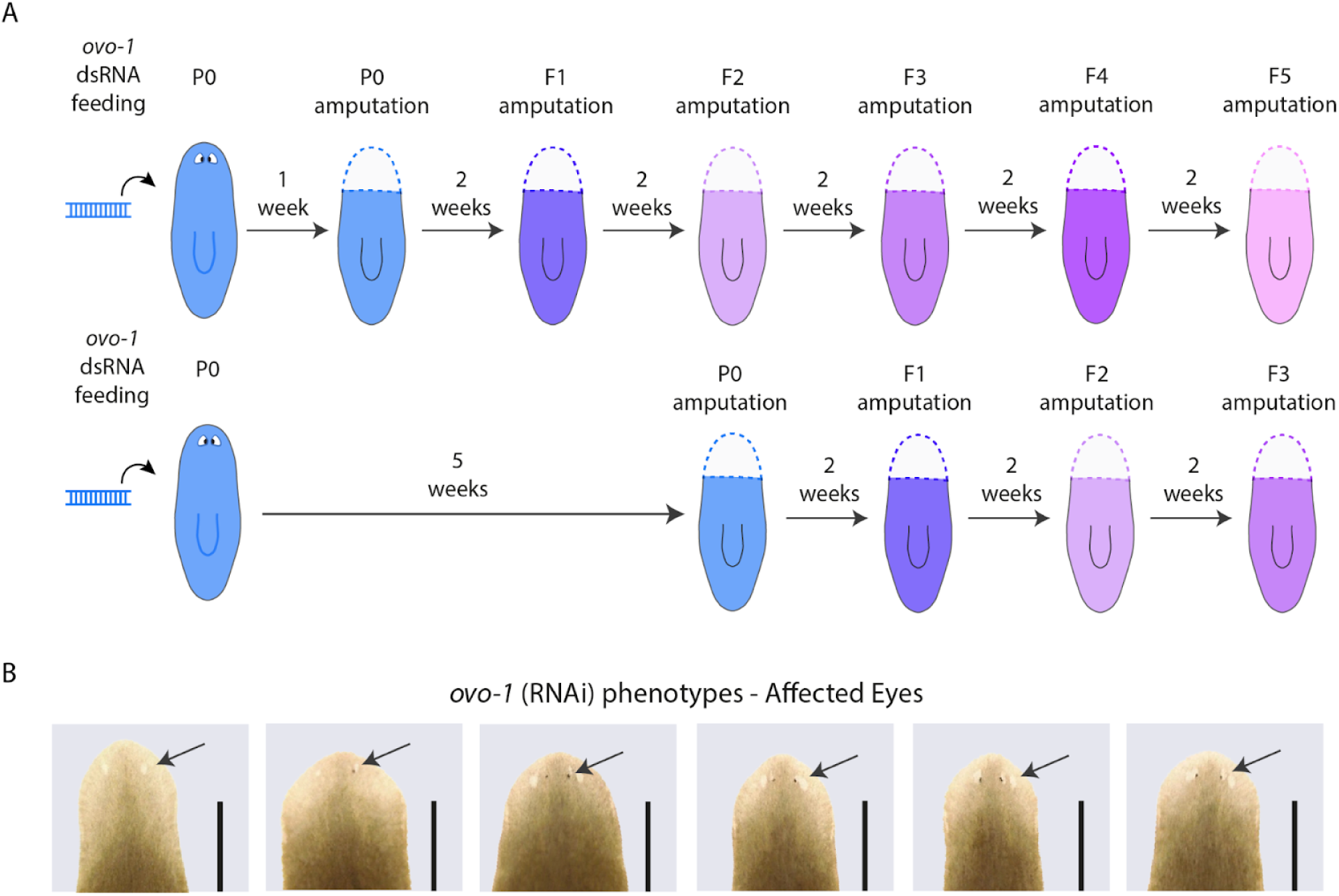
Phenotype persistence is unaffected by the number of amputations in *S. polychroa*. Schematic of the delayed amputation protocol. Standard-cut animals (top) underwent their initial amputation 1 week post-dsRNA feeding, followed by successive amputations every 2 weeks. Delayed-cut animals (bottom) underwent their initial amputation 5 weeks post-feeding, followed by successive amputations every 2 weeks. Distinct colors denote specific amputation-regeneration cycles across both timelines. Representative images defining the “affected eyes” phenotype in *ovo-1* (RNAi) animals. Arrows indicate abnormally regenerated eyes, representing an intermediate, weaker silencing effect compared to the complete loss-of-eye phenotype detailed in Figure 1A. Scale bars = 1 mm.

**Figure S2.**
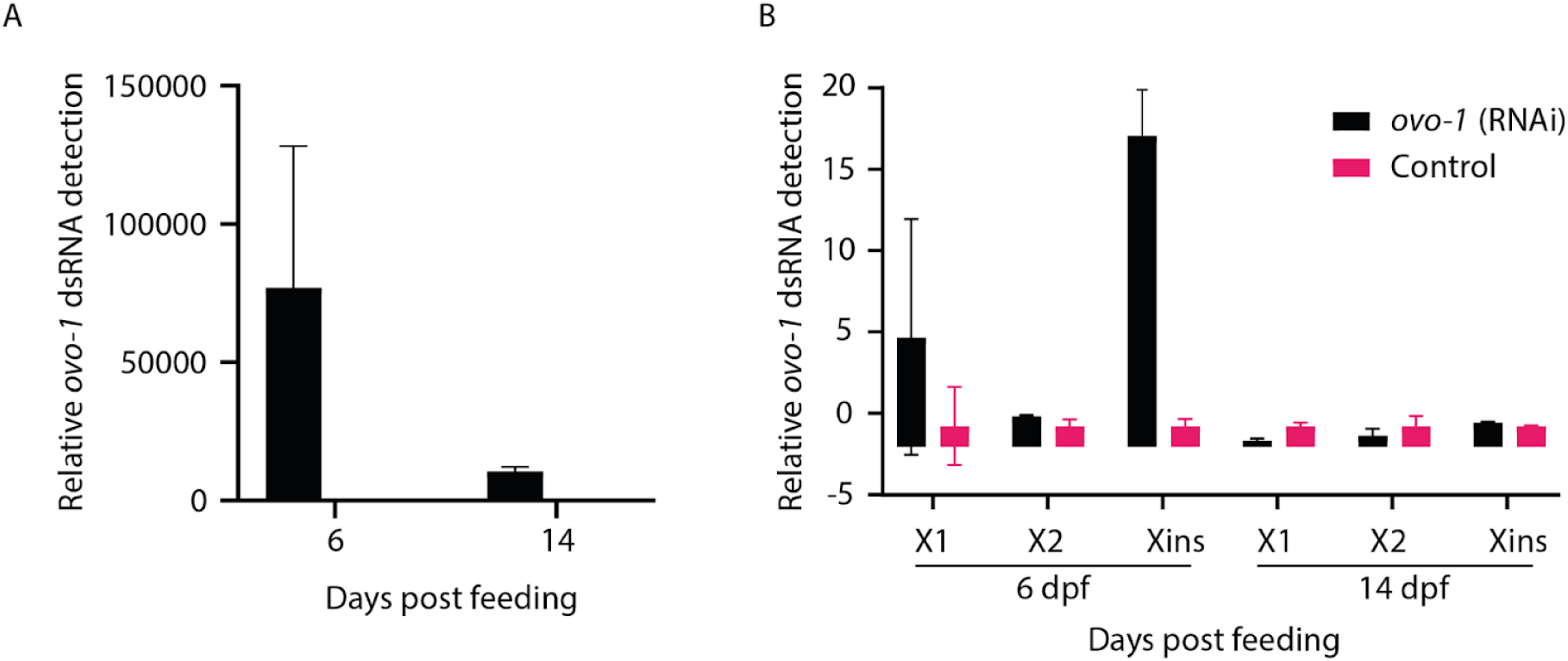
RNAi-mediated gene silencing involves rapid processing of dsRNA. (A) qPCR quantification of *ovo-1* dsRNA in whole-animal extracts. Exogenous dsRNA levels decreased dramatically between 6 and 14 days post dsRNA-feeding (Methods). (B) qPCR quantification of *ovo-1* dsRNA within distinct cell populations fractionated by fluorescence-activated cell sorting (FACS). *ovo-1* (RNAi) and control animals were dissociated at 6 and 14 days post-feeding (dpf) and sorted into neoblast (X1), progenitor (X2), and differentiated cell (Xins) fractions ^41^. By 14 dpf, exogenous dsRNA is nearly undetectable across all isolated cell states. Shown is the average and standard deviation.

**Figure S3.**
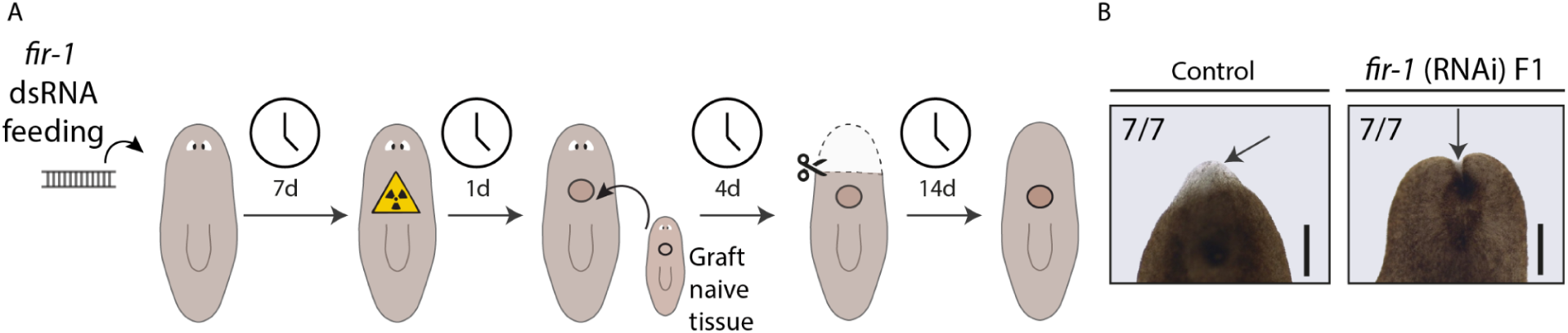
***fir-1* RNAi spreading into naive grafts.** (A) Schematic of the irradiated RNAi host and naive donor grafting assay. *S. polychroa* were fed *fir-1* or control (*unc-22*) dsRNA, then irradiated 1 week post-feeding. Tissue from naive unirradiated donors was then grafted on the Irradiated hosts. After a 4-day integration period, heads were amputated and assessed for blastema formation. Failure to generate blastema indicates the spreading of the RNAi into the neoblasts in the grafted tissue (Methods). (B) Representative images of hosts 6 days post-amputation (dpa). Naive tissue grafted onto *fir-1* (RNAi) hosts failed to form a blastema (arrow, right) and subsequently lysed, demonstrating that the silencing signal spreads from the irradiated host to the naive graft. Control hosts developed normal blastemas (arrow, left). Scale = 1mm.

**Figure S4.**
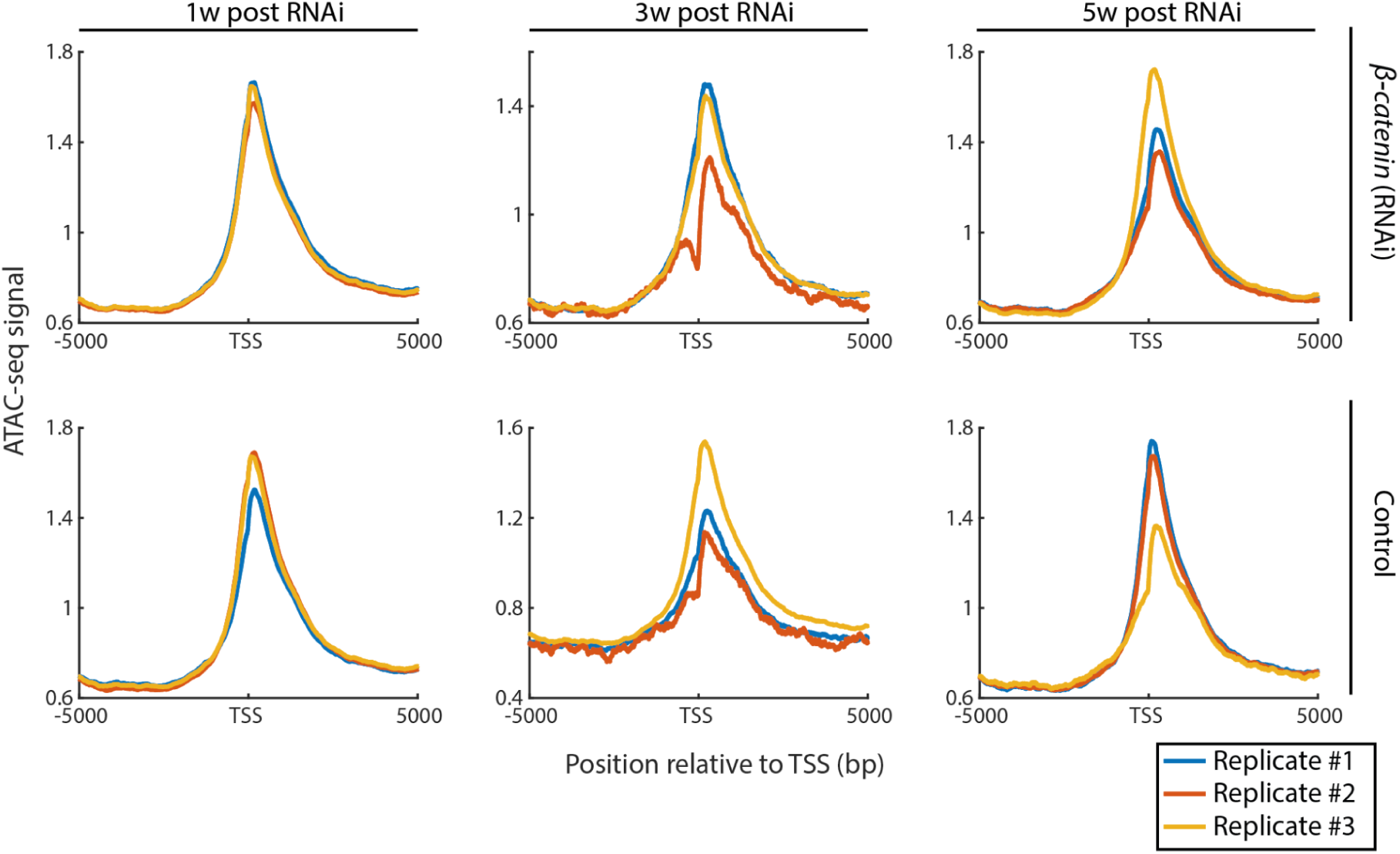
ATAC signal profiles around the transcription start site (TSS) of *Schmidtea mediterranea* genes. *ꞵ-catenin* and control (RNAi) animals were processed for ATAC at different timepoints (1, 3 and 5 weeks) following dsRNA ingestion. In this meta gene analysis, the x-axis shows the genomic position relative to TSS (−5000 to +5000 bp), and the y-axis indicates the averaged ATAC signal intensity.

**Figure S5.**
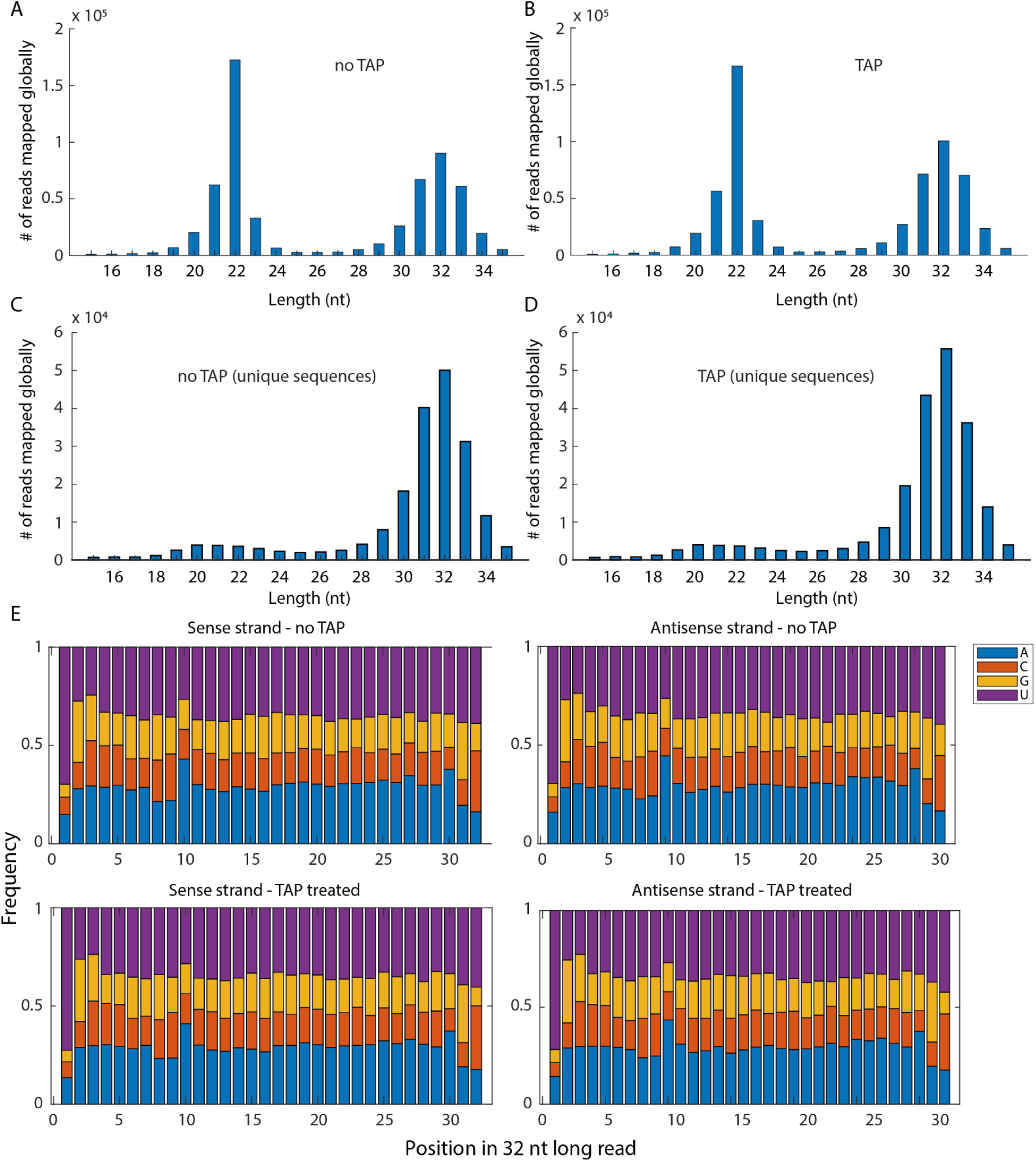
Characteristics and distribution of sRNAs aligned globally to the *S. polychroa* genome. (A, B) Length distribution of all mapped sRNA reads (including redundant sequences) from libraries prepared (A) without or (B) with Tobacco Acid Pyrophosphatase (TAP) treatment. The total read populations exhibit two distinct enrichments at 20–22 nt and 31–33 nt. (C, D) Length distribution of uniquely mapped sRNA sequences from libraries prepared (C) without or (D) with TAP treatment. Excluding redundant sequences reveals that in RNA isolated from naive animals the 31–33 nt species constitute a vastly more diverse population than the 20–22 nt species. (E) Nucleotide frequency across each position of globally mapped 32-nt sRNAs. Sense and antisense reads across all library conditions exhibit a strong preference for uridine at the 5ʹ end (position 1) and an adenosine bias at the 10th position.

**Figure S6.**
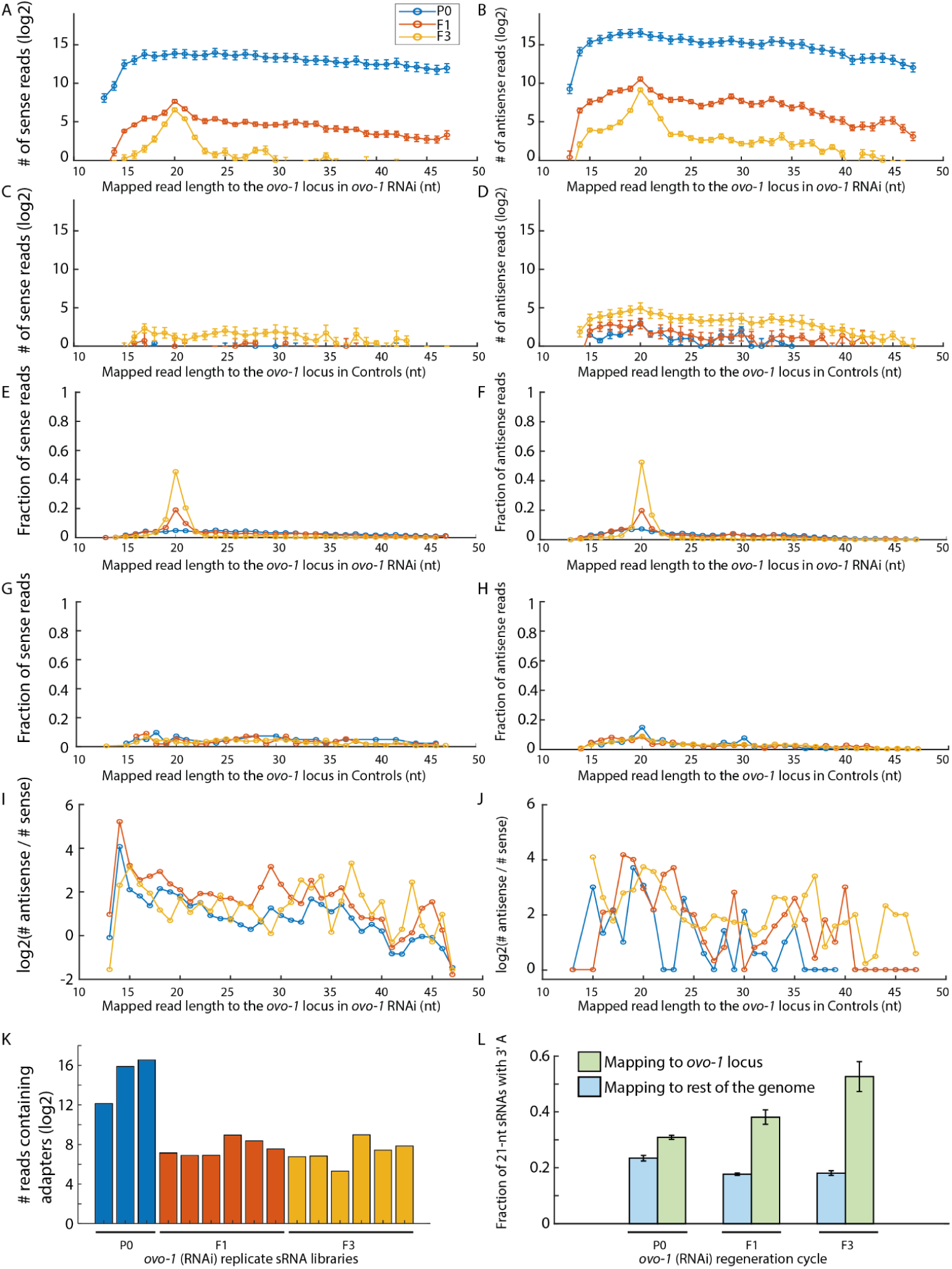
Length distribution and orientation of sRNAs mapping to the *S. polychroa ovo-1* locus. (A–D) Absolute abundance (log2) of uniquely mapped sense (A, C) and antisense (B, D) sRNAs aligning to the *ovo-1* locus following *ovo-1* (RNAi) (A, B) or control RNAi (C, D). (E–H) Fractional distribution of mapped sense (E, G) and antisense (F, H) sRNAs. *ovo-1* (RNAi) animals (E, F) exhibit a pronounced, trans-generational enrichment of 19–22 nt reads in the F1 and F3 generations. Control animals (G, H) have very few reads mapped to *ovo-1* region, and predictably, lack this discrete enrichment. (I, J) Ratio of antisense to sense reads (log2) plotted as a function of read length for *ovo-1* (RNAi) (I) and control (J) animals. (K) Quantification (log2) of reads containing adapter sequences across individual *ovo-1* (RNAi) biological replicates for P0, F1, and F3 generations. (L) Fraction of perfectly mapped 21-nt reads terminating with a 3ʹ adenosine. While the genome-wide mapping does not increase between the initial (P0) and subsequent regeneration time points (F1, F3), the proportion of 3’-A terminated reads mapping to the targeted *ovo-1* locus increases progressively following regeneration. The terminal 3ʹ-A most likely represents non-templated modifications specific to the RNAi response. Bar height represents the average fraction across biological replicates, and error bars indicate the standard deviation.

**Figure S7.**
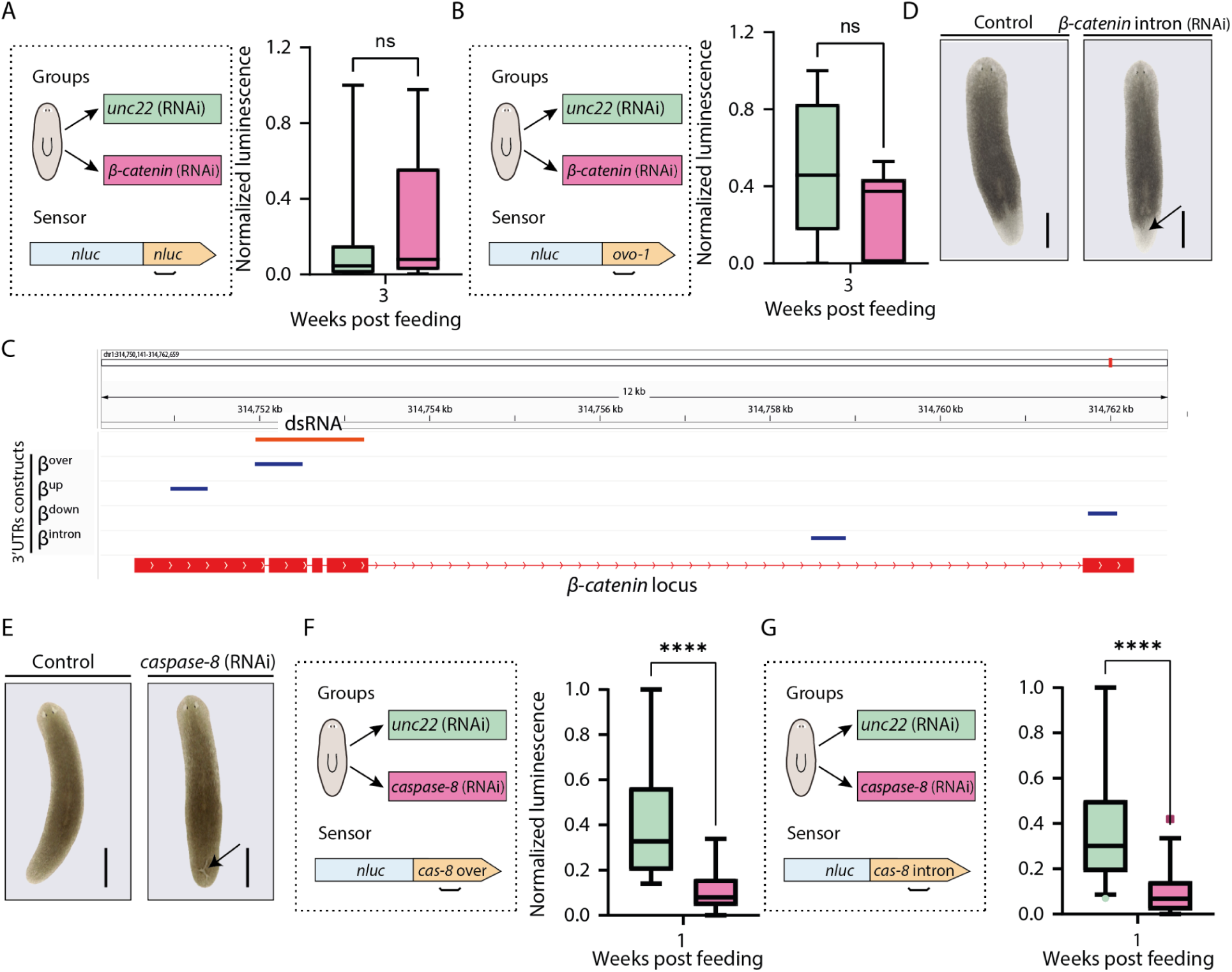
Independent validation of *trans*-silencing spreading using Nluc sensor. (A, B) Specificity controls for the *in vivo* sensor assay. Normalized luminescence of (A) an unmodified *Nluc* sensor and (B) an *ovo-1* sensor at 3 weeks following *ꞵ-catenin* or control RNAi. *ꞵ-catenin* (RNAi) does not suppress the bare Nluc sensor (A) or an off-target sensor (B), confirming the sequence specificity of the sustained silencing effect. (C) Genomic map of the *ꞵ-catenin* locus detailing the exact spatial relationship between the primary dsRNA trigger (thin red block; top) and the specific fragments cloned into the 3’ UTR reporter constructs (thin blue blocks). Exons are shown as thick red blocks. (D) Representative images of animals 14 days post-amputation following two feedings of control or *ꞵ-catenin* intronic dsRNA. Direct targeting of the intronic region fails to induce a regeneration defect (arrow indicates a normally regenerated tail). Scale = 1 mm. (E) Representative images of animals 1 week following two feedings of control or *caspase-8* dsRNA. The arrow indicates a tissue lesion following successful *caspase-8* silencing. Scale = 1mm. (F-G) Trans-silencing dynamics across the *caspase-8* locus. Normalized luminescence of Nluc sensors containing *caspase-8* sequences that (F) overlap the dsRNA target or (G) map to an intron. Both sensors undergo significant suppression 1 week post-feeding, providing independent confirmation that silencing spreads to intronic regions in a separate genetic target. Data are represented as box plots where whiskers indicate minimum and maximum values, horizontal bar indicates median, box shows the interquartile range. For all injection panels, 8-12 animals were injected per group per experiment. Injection experiments were repeated at least twice. Statistical significance was determined using two-tailed Mann-Whitney U tests (ns = not significant; *P < 0.05; **P < 0.01; ***P < 0.001; ****P < 0.0001).

## Notes

### Competing Interest Statement

The authors have declared no competing interest.

